# The grape MYB24 mediates the coordination of light-induced terpene and flavonol accumulation in response to berry anthocyanin sunscreen depletion

**DOI:** 10.1101/2021.12.16.472692

**Authors:** Zhang Chen, Dai Zhanwu, Ferrier Thilia, Orduña Luis, Santiago Antonio, Peris Arnau, Wong Darren, Kappel Christian, Savoi Stefania, Loyola Rodrigo, Amato Alessandra, Kozak Bartosz, Li Miaomiao, Carrasco David, Meyer Carlos, Espinoza Carmen, Hilbert Ghislaine, Figueroa-Balderas Rosa, Cantu Dario, Arroyo Rosa, Arce-Johnson Patricio, Claudel Patricia, Duchêne Eric, Huang Shao-shan Carol, Castellarin Simone Diego, Tornielli Giovanni Battista, Barrieu Francois, Matus J. Tomás

**Author notes:** shared corresponding authorship. shared first-authorship (Zhang Chen, Dai Zhanwu, Ferrier Thilia).

## Abstract

The presence of naturally-occurring color mutants in plants has permitted the identification of many regulatory genes implicated in the synthesis of discrete metabolic compounds, mostly anthocyanins and carotenoids. Conversely, transcription factors that coordinate more than one specialized metabolic pathway seem challenging to screen from a forward genetics’ perspective. We explored the relationship between different branches of the phenylpropanoid and isoprenoid pathways while examining an infrequent berry skin color variegation in grapevine. Red and white berry skin sections were compared at the genetic, transcriptomic and metabolomic levels showing that, as in most cultivated white grape varieties, the uncolored skin section convened the non-functional alleles of the anthocyanin regulators *MYBA1* and *MYBA2*, explaining the lack of pigments. In contrast, light-responsive flavonols and monoterpenes increased in anthocyanin-depleted areas. We disclosed an enrichment of the flavonol, terpene and carotenoid pathways among up-regulated genes from white-skin sections, accompanied by increased expressions of flavonol regulators and the still uncharacterized *MYB24* gene. We used DAP-seq to examine the *in vitro* binding of affinity-purified MYB24 protein to genomic DNA and demonstrated its binding in the promoter regions of terpene (22) and carotenoid (6) genes, in addition to more than 30 photosynthesis and light-response genes, including the flavonol-regulator HY5 homologue (HYH). We confirmed the activation of *TPS35* and *HYH* promoter:luciferase reporters in the presence of MYB24 and the grape bHLH MYC2, all of which correlate in their higher expression in white skin variegated sections. The integration of several datasets allowed to define a list of high confidence targets, suggesting MYB24 as a modulator of light responses including the synthesis of flavonoids (flavonols) and isoprenoids (terpenes, and putatively carotenoids). The correspondence between MYB24 and monoterpenes in all conditions surveyed implies that this regulatory network is broadly triggered towards berry ripening, and that the absence of anthocyanin sunscreens accelerates its activation most likely in a dose-dependent manner due to increased radiation exposure.

## Introduction

Many secondary metabolites provide pigment attributes in plants. Most of them accumulate as the result of the activity of the flavonoid (anthocyanin) and isoprenoid (carotenoid) pathways, two of the most studied specialized metabolisms in plants. Partly, their presence in flowers and fruits has allowed plant co-evolution with insects and seed dispersers, providing a plethora of color hues and tones found in nature. Beyond this purpose, these compounds also filter harmful excessive radiation and provide accessory photosynthetic capacity, among other roles.

As pigments, anthocyanins and carotenoids can be used as reliable markers in forward genetics approaches for the identification of genes underlying their synthesis. Based on this advantage, the most extensive progress in the control of flavonoid pigment synthesis has come from the study of the combinatorial interaction between activating and repressive sets of R3- and R2R3-MYBs, beta helix-loop-helix (bHLH) and trypthophan-aspartic acid repeat (WDR) regulators (1). WRKY transcription factors have been recently shown to interact with these components (2). This ‘MBW-W’ complex (3) is essential for the accumulation of anthocyanins and the acidification of vacuoles, which promote color changes due to the oxidation/reduction of anthocyanin hydroxyl groups.

Besides phenylpropanoid-regulating MYBs, a few isoprenoids MYB regulators have been identified to date, in *Mimulus lewisii* (4), kiwifruit (5) and tomato (6). The overexpression of mimulus RCP1 in a reduced carotenoid pigmentation (*rcp1*) mutant background restored the production of these isoprenoid pigments and surprisingly decreased anthocyanin production by down-regulating the expression of *PELAN*, a MYB activator of anthocyanin biosynthesis. This opposite relation between anthocyanins and isoprenoids has been observed on very few occasions and in some cases with contradictory verdicts (e.g. in tomato) (7). In the case of grapevine (*Vitis vinifera* L.), white grape varieties seem to have higher carotenoid contents compared to dark-skinned cultivars that accumulate anthocyanins in their berry skins (8). Also, terpenes (a group of volatile isoprenoids giving rich aromas) such as linalool are higher in pink-skinned fruits compared to dark red or black cultivars (9). Very recently, it was reported that sesquiterpenes have an antagonistic effect on the accumulation of anthocyanins (10). Altogether, different balancing or opposite relations between anthocyanins and certain isoprenoids seem to exist in the grape berry. Whether this relation depends on transcriptional regulation remains uncertain.

Grapes constitute a rich source of specialized metabolites and thus represent an interesting model to study the connection between different metabolic pathways, particularly as they are all quantitatively influenced by the environment. In the case of pigmented cultivars, anthocyanins begin to accumulate at the onset of ripening (i.e., the widely-used French term *veraison*) in epidermal and subepidermal cell layers that constitute the berry skin (corresponding to L1 and L2 cell types), or also in the flesh (L2) in the case of ‘*teinturier’* cultivars. Their accumulation heightens with light and ultraviolet radiation (11, 12), but declines with high temperatures (13). Mono- and sesquiterpenes vary greatly amongst grape cultivars although a vast majority accumulate at the end of the ripening stage, being highly influenced by temperature (14), light (15), UV-B (16) and water deficit (17). Finally, carotenoids decline progressively through berry skin development (a sharp decrease occurs at the onset of ripening - *veraison*) (18), but this tendency is modified by high radiation levels (19).

The activation of the anthocyanin pathway in the fruit of grapevine depends on the allelic condition of the R2R3-MYBA1 and MYBA2 regulators located on the berry color locus (20). One of the most frequent white-skin phenotypes results from the insertion of the Gret1 retrotransposon in the 5’UTR of *MYBA1* (21) and the concomitant non-synonymous single-nucleotide mutations in the *MYBA2* coding region (20). Due to its transposable nature, the insertion/excision of Gret1 can often occur, leading to somatic mutations in a single cell. If these occur and proliferate in meristematic tissue, pigment mutants can arise as bud sports. Since vegetative propagation is a widely used strategy for grapes, sports and their novel traits can be selected and retained by breeders. In addition, if somatic mutations occur in restricted cell lineages, variegation phenotypes can be observed (22). Berry color depletion and reversion are often observed in vineyards. However, mosaicism or chimerism (i.e. forms of variegation) are somehow more rare events in grapevine and their study is scarce.

Here, we describe the occurrence of a natural berry color variegation found in the black-skinned ‘Béquignol Noir’ cultivar. Red and white berry skin sections were compared to understand the origin and consequences of this color alteration. The use of a likely isogenic background (the only difference being the allelic composition at the berry color locus) provides fundamental new information about the cross-regulation of phenylpropanoids and isoprenoids in response to pigment depletion and establishes a transcriptional association between these two different specialized metabolic pathways in plants. We show that variegation transcriptionally activates the accumulation of a battery of different specialized metabolites that could both filter radiation and/or control oxidative damage. Our results point out MYB24 as an important modulator of this response.

## Results

### Variegated berries lose the capacity to accumulate anthocyanins in the white skin sections due to the inactivation of MYBA1/MYBA2

The grape cultivar (cv.) ‘Béquignol’ is found in Bordeaux and southwest regions of France, with a high predisposition for producing bud sports. This varietal group is composed of three recognized somatic variants; the red/black-skinned cv. ‘Béquignol Noir’, the pale colored cv. ‘Béquignol Gris’ and the unpigmented ‘Béquignol Blanc (Supplementary Figure S1A). In addition to these variants, cv. ‘Béquignol Noir’ vines present in some of their berries an infrequent pigment alteration, occurring in around 53% of the clusters and in about 4% of the berries within those clusters. This variegation phenotype has been observed with persistence over all seasons surveyed for at least 67 years. Berries exhibit uneven skin pigmentation throughout all ripening stages (Supplementary Figure S1B), with small-to-large stripes of white or red color or in some other cases, half-colored berries are also found (Figure 1A).

**Figure 1.**
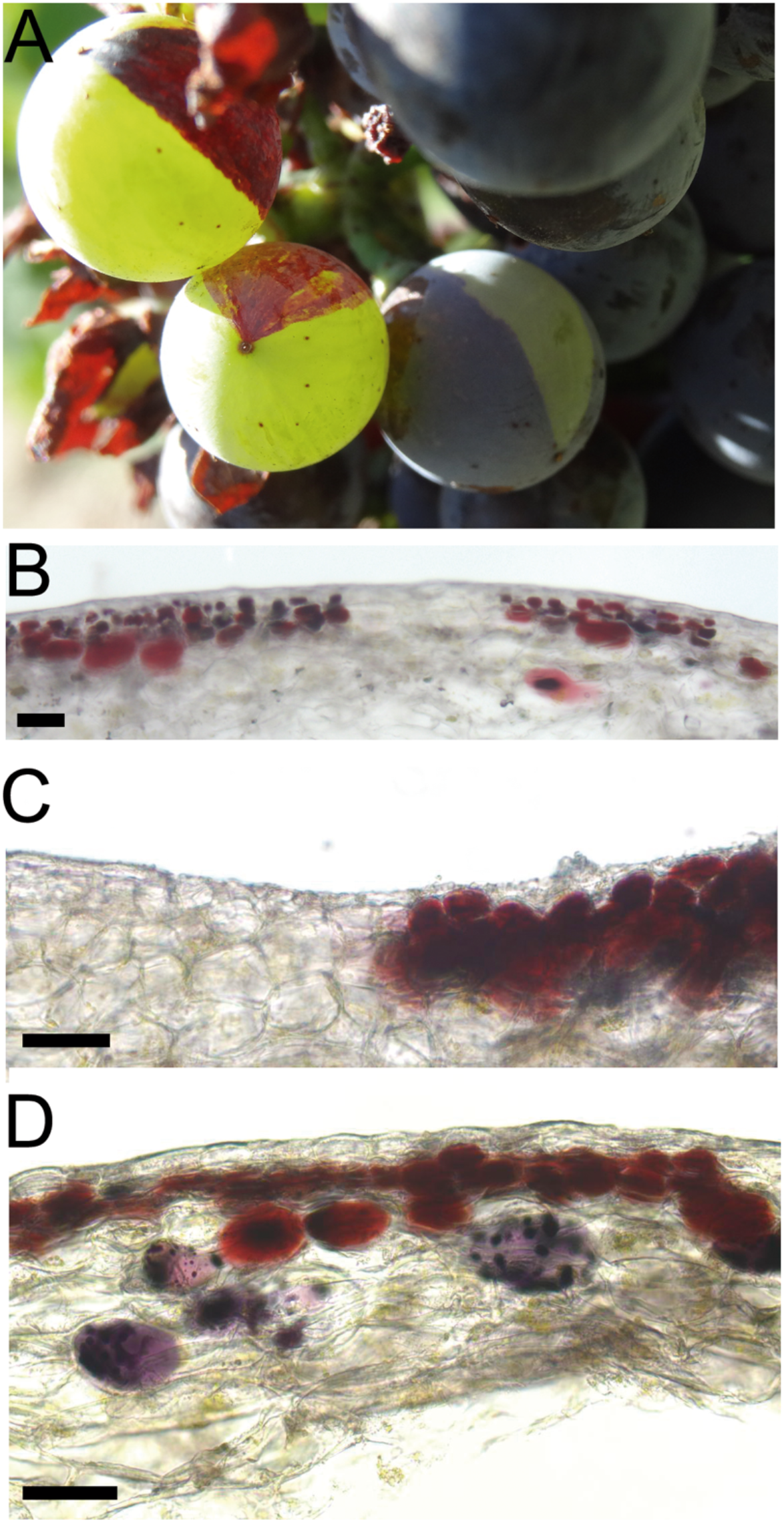
Differential skin pigmentation as a result of anthocyanin depletion in variegated cv. ‘Béquignol Noir’ berries. (A) Irregular berry skin pigmentation observed in field-grown vines at two weeks after veraison (2WAV). (B-D) Light microscopy images of variegated berry skin cell layers. (C-D) Sub-epidermal cells nearby color transitions show anthocyanin vacuolar inclusions (AVIs) accumulating either reddish or purplish pigments corresponding to di- and tri-hydroxylated anthocyanin derivatives, respectively (see also Figure S2A). Pigment accumulation in variegated berries seen throughout fruit ripening, and its comparison to non-variegated cv. ‘Béquignol Noir’, ‘B. Gris and ‘B. Blanc’ fruits can be seen in Supplementary Figure S1. Bar scale: 5μm.

Transversal sections of variegated berries, sampled at 5 weeks after the onset of ripening/veraison (5WAV) were compared by optical microscopy (Figure 1B-D). As seen in both cv. ‘B. Noir’ unvariegated and pigmented variegated berries, skins are constituted by several layers of anthocyanin-accumulating cells. Large and vacuolated subepidermal cells accumulating both reddish and purplish anthocyanin vacuolar inclusion (AVIs) were discernable in pigmented sections. In contrast, anthocyanins were absent in L1 and L2 cell layers of the white skin sections of unvariegated berries. In the variegated berries, the boundaries between pigmented and unpigmented areas were distinguishable. As surveyed by HPLC quantification, anthocyanin derivatives were only present in pigmented berry skins. These were similar in total abundance and relative proportion of di/tri-hydroxylated forms in both unvariegated berries of cv. ‘B. Noir’ and the red skin sections of variegated berries (minor abundance changes were affected by vintage despite proportions of each derivative were not affected; Supplementary Figure S2). The proportion of glucoside derivatives and their coumaroylated and acylated modifications was similar to those previously described for other grape cultivars (23).

As the color of the grape berry skin relies on the allelic condition of a major locus on chromosome 2 that harbors the anthocyanin-promoting R2R3-MYBA1 and MYBA2 transcription factors, we inspected if they could explain this variegation phenotype. The inactivation of *MYBA1* through the insertion of the Gret1 retrotransposon in the promoter/5’UTR region (21) and non-synonymous single-nucleotide polymorphisms in the *MYBA2* coding region (20) account for the un-pigmented phenotype of most white-skinned cultivars known to date. Reversions from non-functional-to-functional alleles largely occur by excision of Gret1 in the form of somatic mutations that occur independently in L1 or L2 layers of floral meristems within buds. Since different plant organs and tissues are derived from the L1 and L2 meristem layers, we followed a cell layer-specific molecular characterization of the berry color locus by assessing eleven molecular markers in genomic DNA extracted from L1+L2 (berry skins and leaves) and L2 (berry pulp and roots)-derived organs from all somatic variants of the ‘Béquignol’ family (Figure 2A). Because ‘B. Gris’ derivates from ‘B. Noir’, the genetic arrangement observed of the berry color locus in ‘B. Gris’ reveals that the only configuration possible for ‘B. Noir’ is the heterozygous (hz) state in the L2 layer. Additionally, this suggests only two possible scenarios of ‘B. Noir’ L1 cell layer: homozygous (hm) or hz for the functional allele. The white skin layer of the variegated berry shows a hm haplotype for the white allele, corroborating the null expression of *MYBA1*, and its target *UFGT1* (Figure 2B). Despite this genetic makeup is identical to ‘B. blanc’, the variegated-red skin area matches the genetic profile of non-variegated ‘B. Noir’ berries. Considering the two hypothetical scenarios of ‘B. Noir’, and the fact that this phenotype is a quite rare event occurring unevenly in the berry, we hypothesize that a somatic recombination event occurred in one L2 cell somewhere before the position of MM SC8_010 and VVNTM4, including both *MYBA1* and *MYBA2* loci. This cell probably originated in the L2 layer with subsequent incorporation into the L1 layer, providing a ‘patch’ presence of non-functional loci. Depending on the berry, one or multiple white patches are observed, suggesting different somatic recombination events taking place.

**Figure 2.**
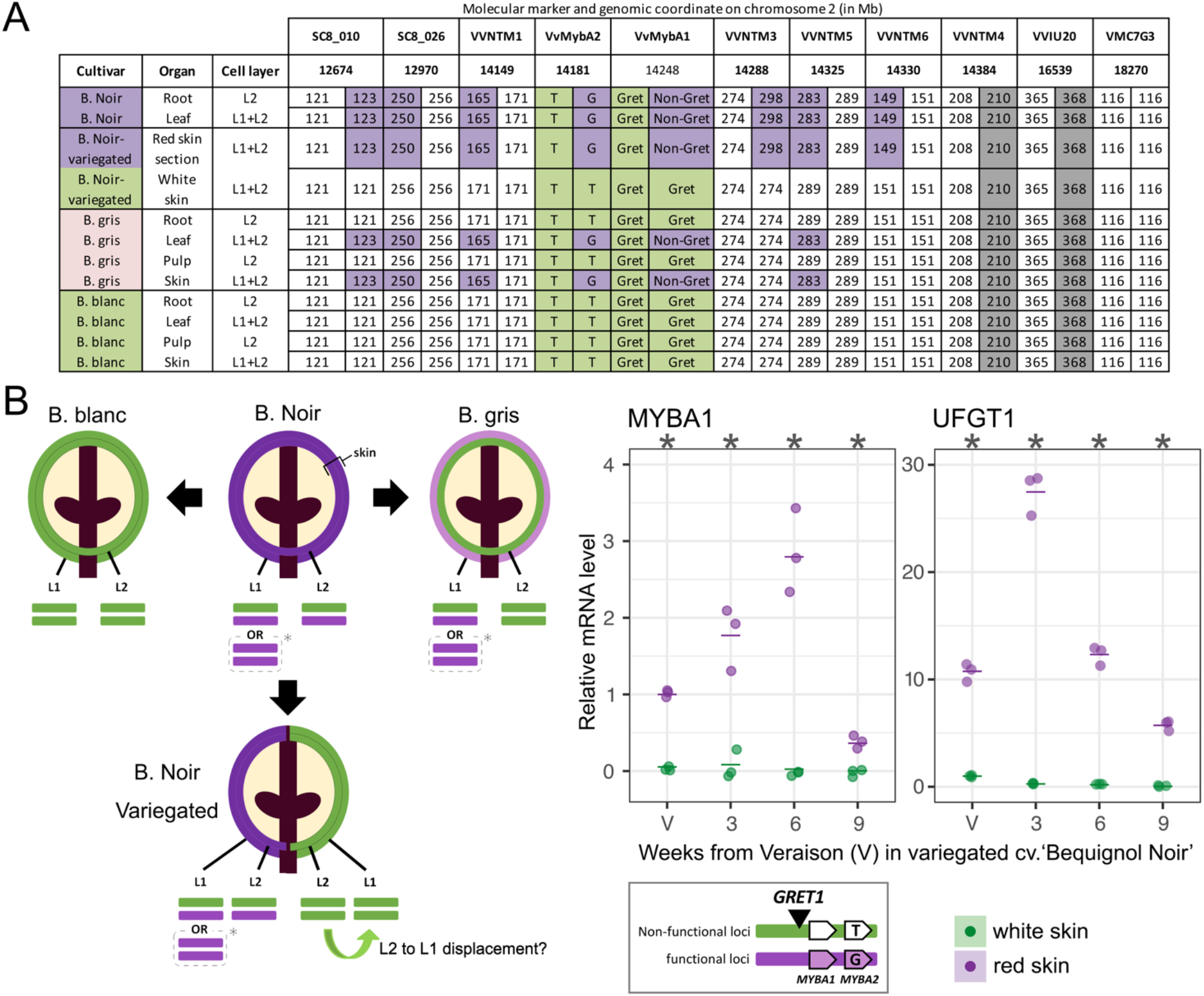
Haplotype structure analysis of the berry color locus show two different genotypes in a single variegated berry of ‘Béquignol Noir’. (A) Genetic profiling of the berry color locus and its surrounding genomic region in color somatic variants, including white, grey, red and variegated berries of ‘Béquignol’ cultivars. Eight microsatellite markers were assessed across the distal arm of chromosome 2, including the allele sequence analysis of the anthocyanin regulators MYBA1 and MYBA2 (allele sizes are shown in base pairs). MYBA1–Gret1 and MYBA2–T are the non-functional alleles, while MYBA1–Non-Gret1 and MYBA2––G correspond to the functional alleles. (B, left panel) Model for the formation of ‘B. Blanc’, ‘B. Gris’, and the variegated phenotype from independent somatic mutation events in ‘B. Noir’, in resemblance to the ‘Pinot’ model (24). The structural dynamics at the berry color locus in the L1 and L2 meristematic cell layers are indicated for each variant. Asterisks represent an alternative configuration for both alleles in the L1 layer. Right panel: expression profiles of *MYBA1* and its target *UFGT1* at different ripening stages in red and white skin sections of variegated berries. About 16 berries from 8 clusters belonging to 5 plants were used per sample. Data from three biological replicates are shown (averages as horizontal lines). Asterisks indicate significant differences (*p*<0.05) between tissues based on one-way ANOVA followed by Tukey’s post hoc test (performed independently for each stage).

### Anthocyanin depletion associates with expression changes in phenylpropanoid and isoprenoid pathway genes and several R2R3-MYB transcriptional regulators

The transcriptomes of the variegated red and white skin sections of cv. ‘Bequignol Noir’ were contrasted using Operon oligonucleotide microarrays (with around 15,000 genes being represented, supplementary Table S1A). The analysis was conducted on grape skins at the mid-ripening stage of 5WAV (Supplementary Table S1B) and showed 807 genes being significantly and differentially expressed by anthocyanin depletion (Supplementary Table S1C), including 454 red-skin up-regulated genes (RUGs, Supplementary Table S1D; from which 301 presented a fold change ≥ 1.5, Supplementary Table S1E) and 353 white-skin up-regulated genes (WUGs, all with fold change ≥ 1.5, Supplementary Table S1F). Gene set enrichment analysis (GSEA, Supplementary Table S1G, H and I) and category enrichment analysis (Mapman Wilcoxon test p<0.01, Supplementary Table S1J) showed that RUGs were enriched in phenylpropanoid pathway terms, including lignin and flavonoid (anthocyanin) biosynthesis and response to heat processes, meanwhile many photosynthesis-related terms (e.g., light reaction, photosystem I and II, chlorophyll metabolic processes) and the carotenoid pathway were significantly enriched within WUGs. WUGs were also associated with other isoprenoids (e.g., terpenes, gibberellins) and response to abiotic stimulus (e.g., light and radiation) (Figure 3A and B, Supplementary Figure S3). In the RUGs and WUGs lists, several differentially-expressed transcription factors appeared, mostly belonging to the R2R3-MYB family. MYBA1 and the shikimate and stilbene pathway regulator MYB15 (25) were induced in red skin areas while the flavonol-regulator MYBF1 (26), the stomata-opening regulator MYB60 (27), and a still-uncharacterized MYB24-homologue were induced in anthocyanin-devoid sections (Figure 3C).

**Figure 3.**
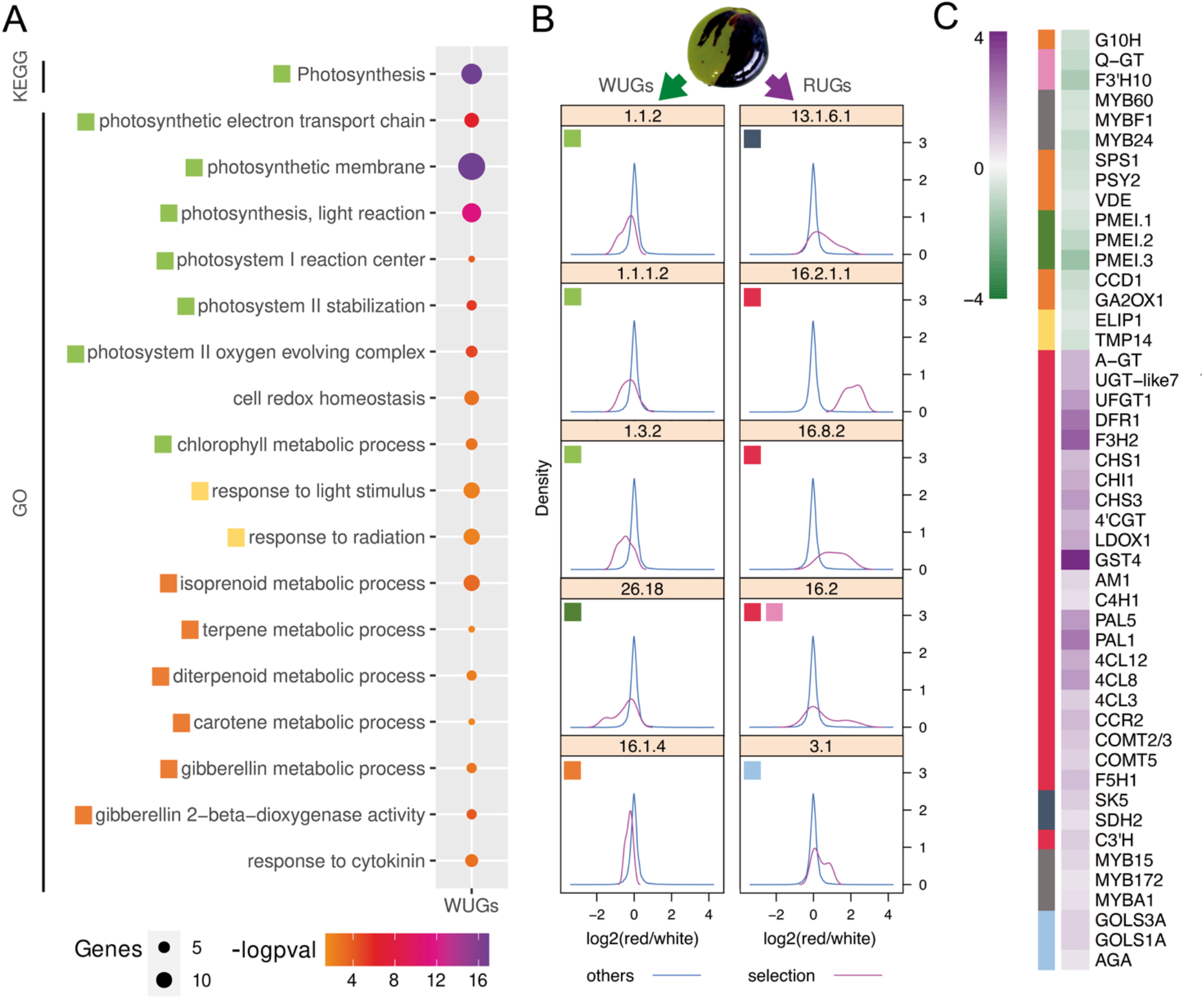
Transcriptomic landscapes of inversely pigmented sections of variegated berries show different pathways perturbed by anthocyanin depletion. (A) A selection of significantly enriched terms in white-skin up-regulated genes (WUGs; complete gene and term lists in Supplementary Table S1F and S1I). (B) Density plots of functional MapMan ontologies in berry skin transcriptomes illustrating shifts in differential gene expression. Significantly affected categories (bins) were determined based on Wilcoxon test (FDR<0.05). Bins: 1.1.2 light reaction, photosystem I; 1.1.1.2 light reaction, photosystem II (polypeptide subunits); 1.3.2 Calvin cycle, rubisco small subunit; 26.18 invertase/pectin methylesterase inhibitor family protein (misc.); 16.1.4 secondary metabolism, isoprenoids, carotenoids; 13.1.6.1 aromatic amino acids metabolism, chorismate; 16.2.1.1 lignin biosynthesis, PAL; 16.8.2 flavonoids, chalcones; 16.2 secondary metabolism, phenylpropanoids; 3.1 minor CHO metabolism, raffinose family. Other significant categories can be found in Supplementary Table S1. (C) Log2 expression ratios of selected genes belonging to significant categories illustrated by colored boxes found in (A) and (B). Gene IDs and expression values are found in Supplementary Table S1K.

### MYB24 expression correlates with several members of the terpene synthase family

Taking advantage of the large amount of public transcriptomic data being available for *Vitis sp*. and the gene co-expression analyses generated with this data (25) and presented in the Vitis Visualization (VitViz) platform (https://tomsbiolab.com/vitviz) (28), we explored the potential gene regulatory mechanisms of the few transcription factors differentially expressed among the WUG lists (MYB24, MYBF1 and MYB60). Aggregate gene co-expression networks (GCNs) (25) were constructed with condition-dependent (flower/fruit; 35 SRA studies) and -independent (all organs; 131 SRA studies) data (Supplementary Table S2A), and analyzed by GSEA. In the condition-dependent data, MYB60 GCN was enriched in ‘cell periphery and plasma membrane’ terms but these were not present in the WUGs-GSEA. Instead, both MYBF1 and MYB24 were highly enriched in ‘photosynthesis-related and terpene synthase activity’ terms, respectively, thus overlapping with the enriched terms found in the variegated WUGs list (Supplementary Table S2B). The GCN of MYB24, the only uncharacterized TF from this list, contained several specialized metabolic genes related to phenylpropanoid, benzenoid and terpenoid compounds and also to hormone (e.g., jasmonic acid, gibberellin), and fatty acid metabolic pathway genes. Several functionally characterized genes involved in the synthesis of mono and sesquiterpenes were present. Among these, a very high correlation was particularly found with TPS35 (VIT_12s0134g00030), an *in vitro* characterized monoterpene synthase (29).

The grape MYB24 (VIT_14s0066g01090) is the only R2R3-MYB factor belonging to subgroup 19 in grapevine (30), and is the closest homolog of the *Arabidopsis* AtMYB24, AtMYB21 and AtMYB57, all of them being highly expressed in inflorescences and promoting flower maturation in a developmental regulatory network also involving ARF and bHLH transcription factors (31–33). In concordance with its characterized orthologues, VviMYB24 shows high expression in flowers, increasing towards the late stage of their development (Supplementary Figure S4A). Additionally, it shows an exponential expression in berry ripening stages towards harvest and post-harvest withering. After inspecting public transcriptomic datasets, we identified MYB24 as being highly expressed in berry skins under two environmental stress conditions: UV-B (in a pigmented cultivar) (16) and drought (white-skinned cultivar) (17). Reanalysis of RNA-seq data also allowed us to identify a splicing variant (MYB24.2) that retained its large intron 2 (>3kb), generating a premature stop codon and leading to an incomplete DNA-binding domain protein (it also lacks a putative transactivation domain located in its C-terminal region) (34). Despite the potentially non-functional MYB24.2 presents in general lower expression, both isoforms share related expression trends suggesting transcriptional regulation as a major process for the late-development specific expression in reproductive organs (Supplementary Figure S4B/C).

Both condition-independent (all organs) and -dependent (flower/fruit) MYB24-centered networks showed closest relationships with *TPS35/09/10/04* (Supplementary Table S2A). To ascertain which are the key developmental stages of flowers and fruits driving the strong correlation between MYB24 and terpenoid pathway genes, we first inspected the cv. ‘Corvina’ (red-skinned) expression atlas (35). Coordinated expression of several *TPS* genes and *MYB24* was most evident in mature flowers and late stages of both berry skin and pulp development (Supplementary Figure S5). We also conducted qPCR expression analyses in flower and fruit development stages of both high and low terpene-accumulating white-skin cultivars (cv. ‘Gewürztraminer’ and ‘Viognier’, respectively). *MYB*/*TPS* expressions were higher in cv. ‘Gewürztraminer’, independently of the stage considered. Among all *TPS* genes surveyed, *MYB24* tightly co-expressed with *TPS35*, as well as with the sesquiterpene synthases *TPS10, TPS14* and *TPS07* in both red and white skinned-cultivars (*TPS09* was annotated only in VCost.v3, so it was not possible to find it in the ‘Corvina’ atlas). *MYB24* and *TPS13* correlated only in the high-terpene accumulating cv. ‘Gewürztraminer’ (Supplementary Figure S6).

### MYB24 binds to a set of terpene synthase genes and flavonol regulators, promoting these pathways in response to berry skin anthocyanin depletion

As grape MYB homologues seemed abundantly co-expressed with terpene synthases, we interrogated whether MYB24 transcription factor could potentially regulate these genes. Despite ChIP-seq being a popular approach for transcription factor binding site (TFBS) discovery, it is limited in scale as it depends on antibody availability and quality, and is challenging for lowly expressed proteins such as transcription factors. Instead, we performed DAP-seq that provides a scalable alternative for non-conventional model species where genetic transformation is difficult (36). The MYB24 coding sequence was amplified from cv. ‘Cabernet Sauvignon’ cDNA, cloned in frame with a HALO-tag at its N-terminal region and *in vitro* pulled-down and mixed with a DAP-seq library prepared with genomic DNA of the cv. ‘Cabernet Sauvignon’ 140x genome-sequenced clone CS08 (37).

Based on peak calling and motif discovery, MYB24 DAP-seq general statistics were similar to those previously described for Arabidopsis R2R3-MYBs (38). The number of peaks were quite similar between the two phased haplotype references (6,869 and 6,170 for the CS08 primary (p) and p+haplotig contigs, respectively). *De novo* discovered motifs from sequences under the 600 most-enriched peaks were nearly identical for both analyses showing the core ‘ACCTAAC’ consensus motif (Figure 4A). This motif is consistent with AC-element consensus sequence (ACC[A/T]A[A/C][T/C] or ACC[A/T][A/C/T][A/C/T]), one of the two main groups of target sequences for R2R3-MYBs (39, 40). As described (41), DNA binding domains of subgroup 19 MYBs in Arabidopsis can associate to AC-rich motifs, an interaction that probably requires an arginine residue in R3 helix 3 (42). The AC-element has also been found in nectary gene promoters (TCACCTAA(C/A)) that are bound by a S19 MYB gene (LxS-MYB305) in tobacco (34).

**Figure 4.**
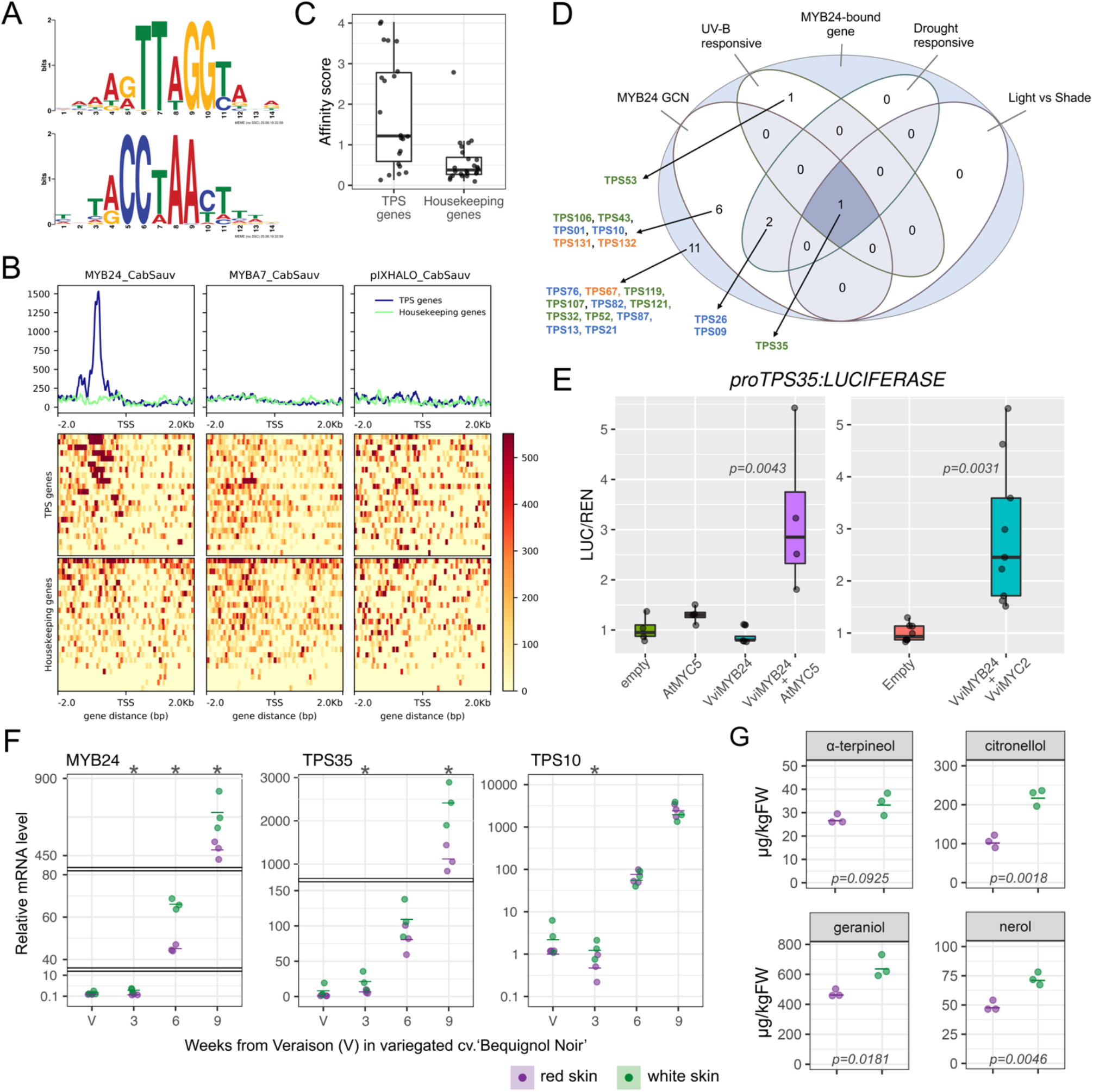
Genome-wide transcription factor binding site (TFBS) discovery by DNA affinity purification sequencing (DAP-seq) identifies TPS35 as a high confidence target of MYB24. (A) Binding motif derived from the 600 most significant peaks of MYB24 DAP-seq, in forward (top) and reverse complement (bottom) directions. (B) DAP-seq binding signal at (−2kb, +2kb) from the TSS (*x* axis) in 22 terpene synthase (*TPS*) genes and compared to background housekeeping genes for MYB24, the anthocyanin regulator MYBA7 and empty vector (pIX-HALO) as negative control. (C) Average motif score (“affinity”) of the MYB24 binding motif across the entire (−2kb, +500bp) region of *TPS* and housekeeping genes. (D) Intersection of *TPS* genes bound and co-expressed with MYB24. (E) Transient expression of VviMYB24 with AtMYC5 (left panel) or VvibHLH07/MYC2 (right panel) activates *VviTPS35* promoter. *Nicotiana benthamiana* plants were agroinfiltrated with 35S:MYB24 and 35S:MYC constructs either alone or in combination (empty vector used as a negative control) and kept in low light conditions for three days before LUCIFERASE activity quantification. Each biological replicate measurement results from the average of two technical replicates. *p* values were calculated based on one-way ANOVA followed by Tukey’s post hoc test. (F) MYB24 and its target *TPS* genes expression profiles at different ripening stages in red and white skin sections of variegated berries of cv. ‘Béquignol Noir’. About 16 berries from 8 clusters belonging to 5 plants were used for each sample. Gene expression data from three biological replicates is shown (averages as horizontal lines). Asterisks indicate significant differences (*p*<0.05) between tissues based on one-way ANOVA followed by Tukey’s post hoc test (performed independently for each developmental stage). (G) GC-MS targeted monoterpene quantifications in red and white skin sections of variegated berries of cv. ‘Béquignol Noir’ at maturity (9WAV). *p* values were calculated based on one-way ANOVA followed by Tukey’s post hoc test.

The 6,170 identified binding sites were assigned to their closest gene feature in the CS08 genome, reaching a total of 5,021 bound genes (Supplementary Table S3A). Peaks were widely distributed throughout upstream, downstream, and inside-gene regions, but a higher proportion was observed close to transcriptional start sites (TSS; Supplementary Figure S7). From all peaks, we found 33 different regions in close proximity to 22 terpene synthase genes belonging to both sesqui- and monoterpene-types (73% at their upstream region, Supplementary Table S3B). *TPS* promoter-preferential binding sites were found specific to this subgroup 19 (S19) member and not, for instance, to Subgroup 6 anthocyanin-promoting MYBs such as MYBA7 (Figure 4B). We searched for the identified AC-element in these 22 *TPS* genomic regions observing an enrichment for a subgroup of *TPS* genes, revealing a preferential binding of MYB24 in *TPS* promoters (Figure 4C). We remapped the DAP-seq reads in the 12X.2 PN40024 reference genome to ascertain the identity of each *TPS*-bound gene, identifying the MYB24 co-expressed *TPS35/09/10/13* genes, among others.

MYB24 is prominently a nuclear-localized transcriptional activator as shown by MYB24:E-GFP localization and Gal4DBD:MYB24 activity assays in tobacco agroinfiltrated cells and yeast, respectively (Supplementary Figure S8A and B). Thus, in order to establish a list of high confidence targets among MYB24-bound genes, we overlapped the DAP-seq data (binding up to −5kb from TSS, Supplementary Table S3C) with MYB24 co-expression data (condition-dependent and independent networks from Supplementary Table S2A, additionally including MYB24-bound *TPS* genes showing MYB24 as part of their own GCN; Supplementary Table S3D) and with reanalyzed transcriptomic datasets from ripening berries where *MYB24* was up-regulated; namely in response to UV-B radiation (NimbleGen microarray data; Supplementary Table S3E) (16), in light versus shade treatments (Illumina RNA-Seq; Supplementary Table S3F) (43) and in response to water deficit (Illumina RNA-Seq; Supplementary Table S3G) (17). Three genes were present in all list sets, including *TPS35*, a putative jasmonate O-methyl transferase (*JOMT*) and a zinc knuckle putative transcriptional regulator (*ZnKn*). Twelve MYB24-bound genes were present in at least three datasets: a UDP-glycosyl transferase of unknown function (*UGT-like1*), a thioesterase, and S-adenosyl-L-methionine:salicylic acid carboxyl methyltransferase (*SAMT*) (Figure 4D, Supplementary Figure S9).

As a high confident target, we tested the transcriptional regulation of *VviTPS35* by VviMYB24 with a dual luciferase assay in tobacco agroinfiltrated leaves. The Arabidopsis AtMYB24 and AtMYB21, which have a role in flower maturation, are required to interact with bHLH factors (AtMYC2/5) to succeed in these roles (32). As expected, the activation of the *VviTPS35* promoter (1.66kb upstream of the ATG) required the co-infiltration with AtMYC5 (Figure 4E). We used AtMYC5’s sequence to search for its homologues in grapevine, identifying the ubiquitously expressed *VviMYC2* (*VvibHLH007*) as the closest (Supplementary Figure S10A and B). The activity of the *VviTPS35* promoter was also dependent on the co-infiltration of VviMYC2 (Figure 4E). Interestingly, VviMYB24 binds in the promoter of VviMYC2 at −6.9kb from the TSS. Further bimolecular fluorescence complementation (BiFC) assay demonstrated VviMYB24 interaction with AtMYC5 and VviMYC2 in *N. benthamiana* agroinfiltrated leaves (Supplementary Figure S10C).

The binding and reciprocal co-expression found with several *TPS* genes (Supplementary Figure S11), together with the direct activation of *TPS35* by MYB24 implied that terpenes should differentially accumulate whenever MYB24 increases its expression, i.e., in late-flower and -berry ripening developmental stages, in white skin sections of cv. ‘Béquignol’ variegated berries and in response to radiation and drought. In fact, this is what we observed; first, high MYB24-terpene correlations were found in flower/fruit development of both high and low terpene-producing white-skinned cultivars (Supplementary Figure S12). Also, by reanalyzing and integrating two metabolomics/transcriptomics studies (17, 44) we found MYB24 to be highly correlated with the accumulation of berry monoterpenes (geraniol, nerol, linalool, and alpha-terpineol) in response to drought (Supplementary Figure S13). We further explored this relationship in the variegated samples by dissecting white and red skin sections, quantifying gene expression throughout all ripening stages and by using targeted and untargeted GC-MS metabolomics at the harvest stage (9WAV). *MYB24, TPS35*/*TPS10* showed very similar expression profiles with an exponential behavior increasing towards the late stages and increasing their expression in the white skin sections in at least one time point (Figure 4F). Among volatile compounds with higher accumulation in white-variegated berry skin sections we identified the monoterpenes citronellol (in both targeted and untargeted assays), geraniol, nerol and alpha-terpineol (Figure 4G, Supplementary Figure S14).

We suggest 418 additional putative MYB24 targets based on the overlap of DAP-seq with at least one of the four considered datasets (mapped to PN40024 12X.2 assembly and associated to its VCost.v3 annotation; Supplementary Table S3H). These include the light-responsive flavonol-pathway regulators HY5 HOMOLOGUE (HYH) and MYBF1 (45) and several photosynthesis-related genes. VviHY5/HYH control several early light responses, including the flavonol accumulation through activation of the regulator MYBF1 and metabolic pathway genes such as flavonol synthase 1 (*FLS1*) and the flavonol glycosyl-transferases *GT5/6* (45, 46). MYB24 binds the HYH promoter at around −0.92kb from the TSS (Figure 5A) and MYBF1 at −3.6kb upstream of the start codon. A dual luciferase assay in tobacco agroinfiltrated leaves demonstrated that the activity of HYH promoter was enhanced by the co-infiltration of VviMYB24 and VviMYC2 (Figure 5B). As expected, *HYH, MYBF1*, and their target *FLS1* and *GT5* were induced in the white skin sections of variegated berries throughout berry ripening (Fig 5C). The flavonol content of white skin sections quantified at 5WAV showed increased levels of the glycosylated forms of quercetin and kaempferol when compared to red skin sections. In particular, the content of two flavonols, quercetin-3-glucoside (q-3-glc) and q-3-glc-6-ac, in the white skin sections of the variegated berries tripled the amount found in pigmented skin samples. Quercetin-3-gal and kaempferol-3-glc were also more abundant in the variegated white skin sections (Figure 5D). Finally, the integration of the two previously-included drought studies also showed a high correlation of MYB24 with kaempferol and quercetin glucosides (Supplementary Figure S13).

**Figure 5.**
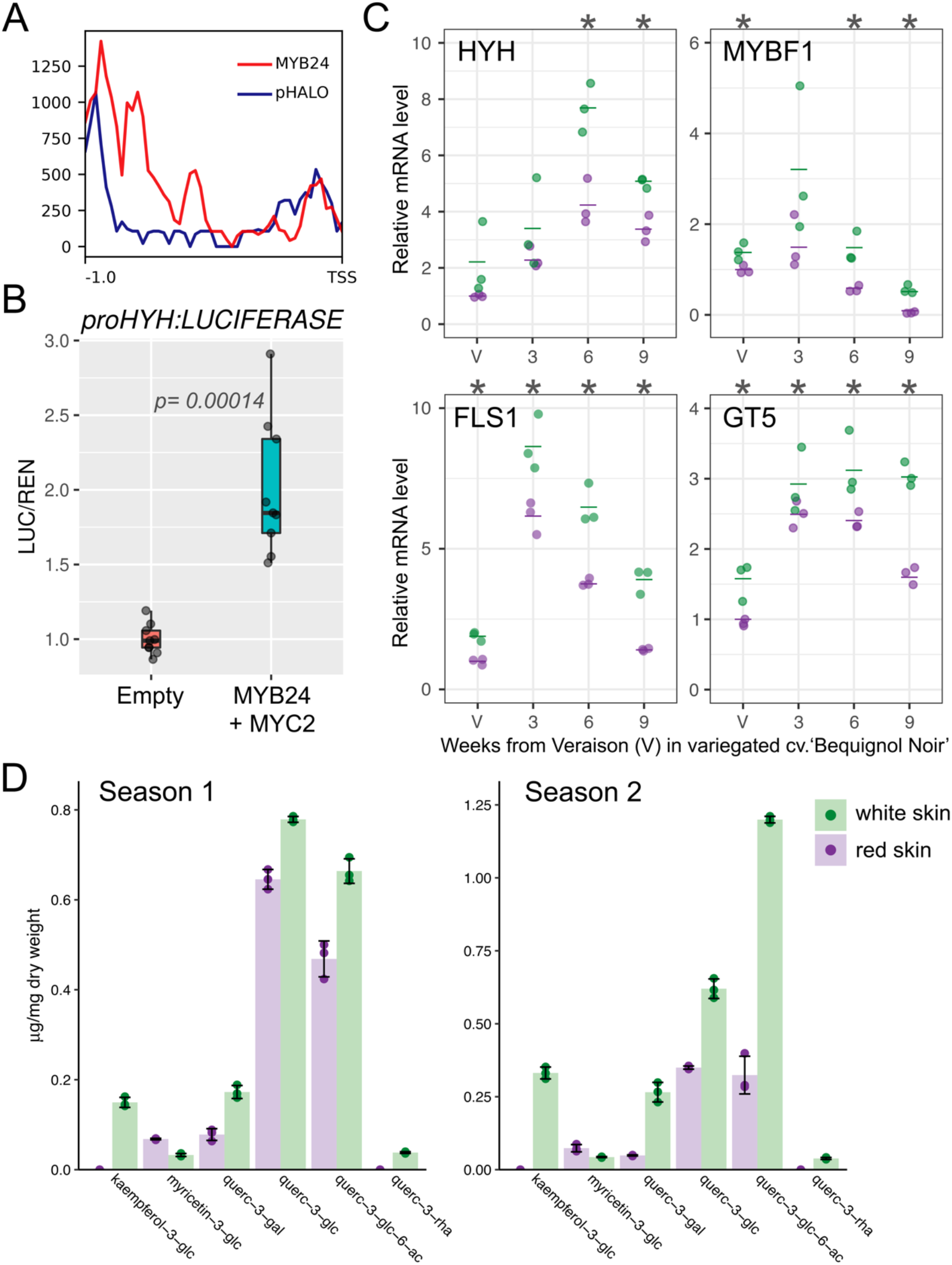
MYB24 promotes flavonol accumulation through binding and regulation of the flavonol-regulator HYH. (A) MYB24 binds to the promoter of the light-early response regulator HY5 homologue (HYH). DAP-seq binding signal at −0.92kb from the TSS (*x* axis), compared to empty vector (pIX-HALO) as negative control. (B) Transient expression of VviMYB24 with VvibHLH07/MYC2 activates *VviHYH* promoter. Each biological replicate measurement results from the average of two technical replicates. *p* values were calculated based on one-way ANOVA followed by Tukey’s post hoc test. (C) The expression profiles of flavonol-related genes (*HYH, MYBF1, FLS1*, and *GT5*) at different ripening stages in red and white skin sections of variegated berries of cv. ‘Béquignol Noir’. About 16 berries from 8 clusters belonging to 5 plants were used for each sample. Gene expression data from three biological replicates is shown (averages as horizontal lines). Asterisks indicate significant differences (p<0.05) between tissues based on one-way ANOVA followed by Tukey’s post hoc test (performed independently for each developmental stage). (D) Flavonol composition in ‘B. Noir’ variegated berry skins at 5WAV, at two consecutive seasons (vintages). High performance liquid chromatography (HPLC) quantifications are expressed as μg/mg of dry weight of quercetin-3-glc equivalents. Standard error bars were calculated from biological replicates.

### *MYB24* expression is modulated by light exposure in a dose-dependent manner

*MYB24* is highly induced by radiation as found in the reanalyzed ‘UV-B responsive’ (16) and ‘light vs shade’ (43) transcriptomic datasets. We confirmed that this light responsiveness occurred throughout all ripening stages by performing qPCR in berry skin samples obtained from light exclusion, UV-B filtering and UV-B irradiance treatments conducted in cv. ‘Cabernet Sauvignon’ field and greenhouse plants (11, 45, 46). In all cases, light and UV-B positively influenced MYB24 expression, a response mirrored by *TPS35* (Supplementary Figure S15) and *HYH* expressions.

The light-responding behavior of *MYB24* drove us to inspect whether anthocyanins, known to filter sunlight in plant tissues, could influence the response of *MYB24* to light, and if this could explain its lower expression in the variegated red skin sections of cv. ‘Béquignol’. We inspected gene expression levels in response to light at two depths within the berry pericarp (i.e., skin and pulp) in the cv. ‘Gamay’ and its ‘*teinturier’* somatic variant cv. ‘Gamay Fréaux’, characterized by the accumulation of anthocyanins in the pulp starting at the onset of ripening (47). As expected, shade drastically reduced the expressions of *MYB24, TPS35* and *HYH* in both tissues in most post-veraison time points surveyed (Figure 6A, Supplementary Figure S16), but additionally, we observed an influence of tissue and cultivar suggesting a positional effect on the expression of *MYB24* that resembles a light dosage response (i.e., the inner pulp tissue being less light responsive than the skin). This effect was more evident at the late stage, when *MYB24* displays its highest expression (Supplementary Figure S16). Furthermore, when considering the levels of anthocyanins in these samples we could observe a clear negative correlation between *MYB24* and the amount of pigments at late ripening stages when independently analyzing skins and pulps (Figure 6B), corroborating this sunscreen effect of anthocyanins. Anthocyanin content would also explain a diminished *MYB24* expression in ‘Gamay Fréaux’ compared to ‘Gamay’.

**Figure 6.**
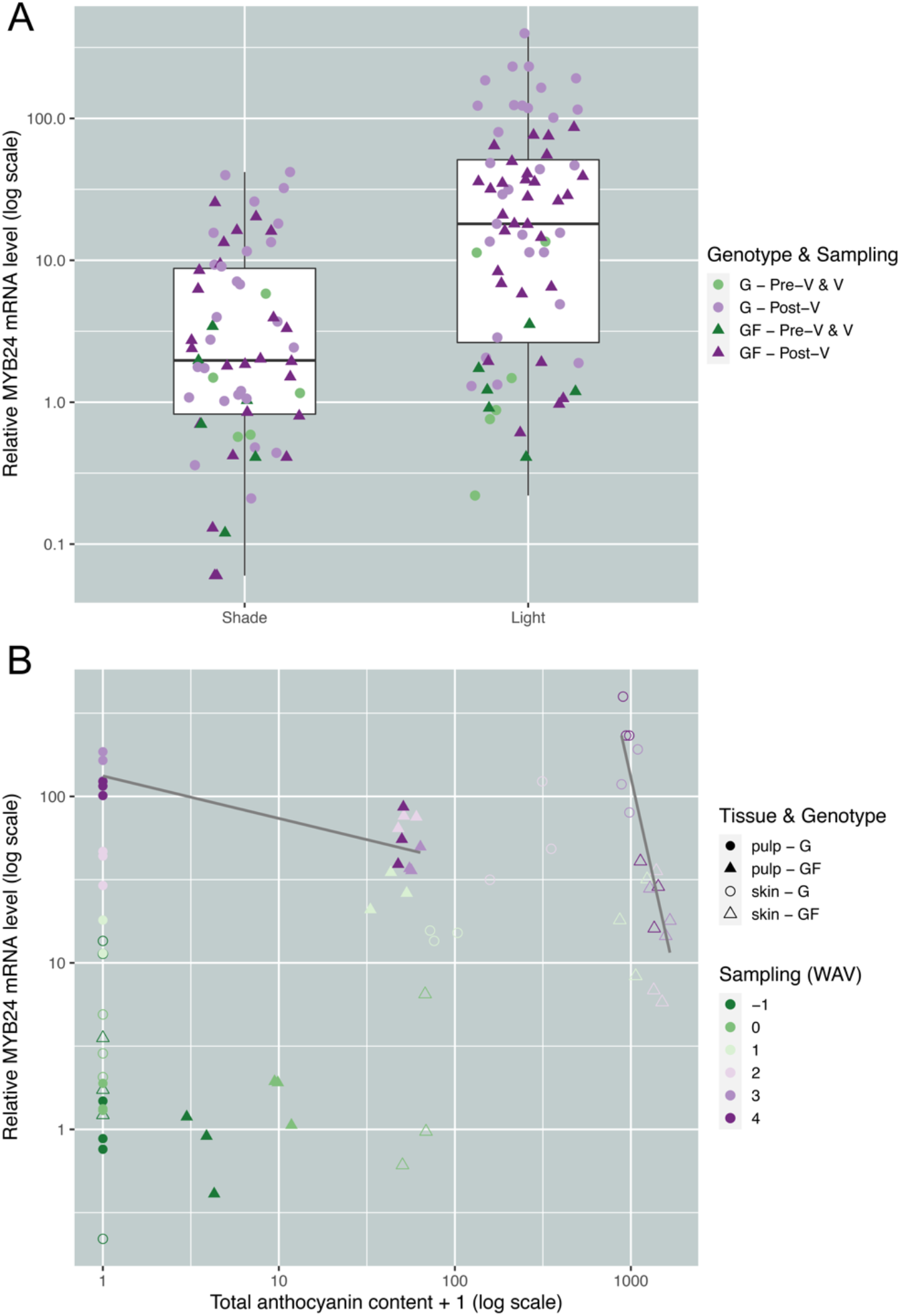
*MYB24* responds to light influenced by anthocyanin berry accumulation. (A-B) *MYB24* expression increases upon developmental stage but its negatively influenced by shade, sunscreen (i.e., anthocyanin) accumulation and berry tissue position. Skin and mesocarp gene expression responses to sunlight exclusion were obtained from field trials of cv. ‘Gamay’ and its ‘*teinturier*’ (red-flesh) somatic variant cv. ‘Gamay Fréaux’. A complete fruit sunlight exclusion treatment was imposed by covering grape clusters with opaque boxes (from two weeks before veraison till maturity) and compared to grape clusters exposed to natural light conditions as control (100% light incidence). Developmental stages correspond to −1 to 4 weeks after veraison (WAV). *MYB24, TPS35 and HYH* gene expression profiles are found in Supplementary Figure S15.

## Discussion

### Anthocyanins, as protective epidermal sunscreens, fail to evenly accumulate in variegated berries leading to the accumulation of photoprotective flavonols and terpenes

Despite light being an absolute requirement for the existence of most life forms on earth, excessive irradiation can generate substantial damage to biological systems, including DNA beaks caused by pyrimidine dimers, reactive oxygen production and reduction in plants photosynthetic capacity by impairments of photosystems (48). In sessile organisms, these deleterious effects have urged the production and selection of an unceasing list of specialized metabolites, which are not only capable of reflecting or filtering light but also of repairing potential harm produced by excessive radiation. For example, anthocyanins probably emerged in plant evolution offering different coping solutions in parallel: protecting from oxidative damage and aiding robust sunscreen capacity. On the other side, anthocyanins also play attraction roles, acting as cues for pollinators and seed dispersers, and were therefore crucial for the co-evolution of animals and angiosperms (49). The accumulation of these compounds in fruits such as those of grapevine allows the plant to interact and protect themselves from their environment.

Anthocyanins are pigments found in the majority of plant species. However, variations in the structural and regulatory genes controlling their accumulation have enabled plant organs to display diverse palettes of colors, hues and tones in nature. In grapevine, berry coloration of red-skinned cultivars begins at the onset of ripening, when anthocyanins start to accumulate in vacuoles and anthocyanin vacuolar inclusions of epidermal and hypodermal cells of the skin. Anthocyanins also accumulate in mesocarp cells in the case of ‘tenturier’ cultivars, some of which have been molecularly associated with duplication of MYBA1 binding sites in its promoter increasing its own expression (50). The *Vitis vinifera* ‘Béquignol Noir’ cultivar is a red-skinned cultivar presenting a rather small proportion of variegated berries with uneven skin pigmentation. This is a stable phenotype that has been consistently recorded at least the last 67 years in a field collection. In this study, we show that anthocyanins are exclusively accumulated in pigmented skin sections of variegated berries, presenting similar derivative diversity and quantity compared to unvariegated ‘B. Noir’ berries, and as quantified in many other red-skinned cultivars (23).

As largely described for many different cultivars and their somatic variants (51–53), depletion of anthocyanins usually results from deletions or mutational-induced inactivation of MYBA1/A2 transcription factors. Two of the most frequent alterations are in the insertion of the Gret1 retrotransposon in the MYBA1 promoter/5’UTR (5’-untranslated region) and single nucleotide substitutions/deletions in MYBA2 coding region (20, 21). As presented here, the cv. ‘Béquignol Noir’ white skins of variegated berries show a genetic configuration resembling that of white cultivars, despite the rest of the plant being heterozygous for *MYBA1*/*A2* functional alleles. Variegated tissues, as a form of mosaicism, are often described as periclinal chimeras, where cells are displaced from different cell layers. Our results suggest the occurrence of a functional inactivation of MYBA1/A2 in layer L2 meristematic cells, some of which have further invaded, gradually and heterogeneously, the epidermal cell layer (L1). This invasion of epidermal L1 cells by L2 unpigmented sub-epidermal cells has been suggested to explain the phenotype of cv. ‘Shalistin’, from cv. ‘Malian’ (54).

Despite variations in anthocyanin accumulation offer a vast and desired diversity of colors in grapevine fruits (many of which are reflected in the wines being produced), the decrease or complete depletion of anthocyanins in the skin epidermis renounces the protective advantages endowed by these pigments. Under this scenario, epidermal layers of berry skins have to design and execute an alternative plan to shelter from the effects of excessive light and ultraviolet radiation. The accumulation of flavonols in response to excessive light represents one of the fastest metabolic responses to environmental stresses ever described in plants (55). This is because flavonols play important roles as antioxidants in photoprotection (56). The increased accumulation of flavonols in the white skins of our study subject allowed us to hypothesize that anthocyanin-devoid sections were responding more intensely to sunlight.

Our results also suggest that isoprenoids, in particular monoterpenes, also form part of the berry’s ‘plan B’ in the response to higher radiation caused by anthocyanin depletion. Potential roles of monoterpenes dealing with oxidative stress have been previously suggested, as to directly mitigate ozone levels and scavenge ROS leading to decreased oxidative damage and improved thermotolerance (57, 58). In fact, the increased content of the monoterpenes terpineol, citronellol, geraniol and nerol increased significantly in the non-pigmented sections of variegated berries. Despite previous studies in different plant species have shown how terpenes increase in response to radiation (e.g., in peach) (59), less is known on how terpene synthases are transcriptionally activated by light. In grapevine, most observations suggest that terpene responses to light depend on cultivar and developmental stage in addition to the influence of additional environmental factors. UV-B filtered fruits showed a decreased expression of monoterpenoid biosynthetic genes (16) and monoterpene metabolic genes were positively influenced by light (60). Also, one of the suggested causes for berry terpene accumulation under drought (17) is the fall of leaves as a direct consequence of water restrictions, leading to increased sun exposure of fruits.

### Anthocyanin depletion drive berries to transcriptionally respond to radiation through a MYB24/HYH module

Our transcriptomics and qPCR gene expression analyses of the variegated berries showed several genes associated with photosynthesis (including photosystem function), carotenoid metabolism and light-induced responses as being significantly induced in white-skin sections, in addition to the regulatory and structural pathway genes related to the accumulation of flavonols and terpenes. The activation of these genes corroborates the idea that white skins are experiencing increased radiation as a result of the lack of anthocyanin sunscreens. In the virtue of exploring how variegated berries may activate terpenoid and flavonol metabolism and other protective mechanisms due to the uneven distribution of skin anthocyanins during ripening, we searched for expression changes in transcription factors potentially governing these responses. Our transcriptomic/metabolomic metanalyses, cistrome data and their validation through several approaches suggest that these changes are governed transcriptionally through the activity of a few transcription factors from the R2R3-MYB (MYB24 and MYBF1) and bZIP families (HYH).

MYB24 binds to several photosynthesis and light response-related genes such as YCF3, OHP2 and ELIP1(61, 62), among many others. The DAP-seq data of MYB24, its expression behavior in terms of light-responsiveness and its high correlation with the increase of light-responsive genes and metabolites in all the datasets generated and reanalyzed, evidences a major role of this TF in the general light-signaling pathway. Gene ontology analysis of MYB24 bound genes showed ‘RNA biosynthesis-Transcriptional regulation’ as a highly significant enriched term; in fact around 7.3% of its target genes encode for transcription factors (Supplementary Table S3I and Supplementary Figure S17), placing MYB24 in a leading hierarchy of transcriptional regulation. Among MYB24 targets being transcription factors, we identified *HYH* and *MYBF1* genes, both coding for TFs largely associated with light and UV-B radiation responses, in grapevine and several other species. HY5 and HYH act as partially redundant central mediators of photomorphogenic responses and are considered as marker genes of light signaling. In *Arabidopsis*, they bind to several photosynthesis and light protection-related genes (63). Because of their ‘very early’ behavior we cannot rule out the possibility of a feedback regulation of MYB24 by any of these bZIP regulators. As shown here, the control of flavonol accumulation by MYB24 seems to be indirect, i.e., through the activation of HYH and MYBF1 as intermediate regulators. Despite this, it seems to be a stable process as MYB24 has been previously identified among several metabolic QTLs segregating with flavonol content in ripe berry skins (64).

### Coordination of isoprenoid and phenylpropanoid metabolisms by MYB24

Previous studies have evidenced an opposite relationship between carotenoid or terpenes (or their related genes) and anthocyanins, especially when comparing cultivars with different degrees of pigmentation (8–10, 65). Our data situate MYB24 and its regulatory network in the center of this conjuncture. Anthocyanin depletion results in excessive irradiation in white-skin sections stimulating the expression of MYB24. On the contrary, skin-localized anthocyanins seem to produce a self-shade effect over the mesocarp cells, reducing MYB24 expression in pulp compared to skins. The inverse correlation of *MYB24* and *TPS* gene expressions with anthocyanin berry skin accumulation at late stages of ripening is even evidenced when comparing pink and dark red cultivars (66).

VviMYB24 forms part of Subgroup 19 (S19) within the R2R3-MYB transcription factor family. This subgroup is mainly involved in the maturation of flower organs, a role potentially conserved in grape due to the high expression of *VviMYB24* in this organ. Flowers are known to produce large amounts of volatile organic compounds (VOCs, including terpenes) which participate in several processes such as pollinator attraction and herbivory defenses. In *Arabidopsis*, S19 mutants show a mis-regulation of several *TPS* genes (33), including *AtTPS03* (the closest homologue of *VviTPS35*). Very recently it was suggested that the jasmonate-regulated AtMYB21 could activate the expression of the terpene synthase genes *AtTPS14*/*21* (67). Following the same behavior, we show that MYB24 binds to several *TPS* genes, preferentially at their 5’ regulatory region, suggesting its capacity to regulate them. The inspection of MYB24 ‘very high confidence genes’ indeed shows an enrichment of the term ‘terpene synthase activity’.

Our results suggest that MYB24 could potentially regulate two additional metabolic pathways. First, MYB24 binds in close proximity within the promoter with at least five genes related to isoprenoid and carotenoid metabolism: the farnesyl diphosphate synthase (FPS), GGPS1, carotenoid isomerase (CRTISO2), 9-cis-epoxycarotenoid dioxygenase (NCED6) and a lycopene epsilon cyclase (LCYE). The clearest role of MYB24 in modulating light responses through carotenoids is observed with the carotenoid isomerase and the lycopene beta cyclase, highly induced in the white skin sections of variegated berries (Supplementary Figure S18A). LCYE produces alpha-carotene from lycopene and is the first committed step in the production of lutein, the most abundant carotenoid (xanthophyll) in photosynthetic plant tissues where it plays important roles in light-harvesting complex-II structure and function (68, 69). In the second place, MYB24 may also regulate the production of volatile benzenoids. In petunia, its homologues EMISSION OF BENZENOIDS I/II (EOBI/II) regulate the expression of eugenol synthases (IGS) and an ABCG1 transporter leading to the accumulation of eugenol/isoeugenol, two VOCs belonging to the phenylpropanoid pathway that is emitted at night as part of the floral scent bouquet of petunias to attract pollinators (70–73). In line with these observations, our DAP-seq data also showed that MYB24 binds to *IGS2* (*VIT_03s0088g00140*) and *ABCG1* (*VIT_03s0017g01280*) in correlation with eugenol being more accumulated in white-skin sections of the variegated berry (Supplementary Figure S18B).

Very few transcription factors are known to control terpene synthesis in model plant species (67, 74, 75) and even less are known as regulators of more than one metabolic pathway. Our study shows that the control of multiple metabolic pathways is possible, in this case by the R2R3-MYB24 transcription factor.

## Acknowledgements

This work was supported by Grant PGC2018-099449-A-I00 and by the Ramón y Cajal program grant RYC-2017-23645, both awarded to J.T.M., and to the FPI scholarship PRE2019-088044 granted to L.O. from the Ministerio de Ciencia, Innovación y Universidades (MCIU, Spain), Agencia Estatal de Investigación (AEI, Spain), and Fondo Europeo de Desarrollo Regional (FEDER, European Union). C.Z. is supported by China Scholarship Council (CSC) no. 201906300087. C.Z. is supported by China Scholarship Council (CSC) no. 201906300087. K.G. and Z.R. were supported by the Slovenian Research Agency (grants P4-0165 and Z7-1888). S.C.H. is partially supported by National Science Foundation grant PGRP IOS-1916804. This article is based upon work from COST Action CA 17111 INTEGRAPE, supported by COST (European Cooperation in Science and Technology). Data has been treated and uploaded in public repositories according to the FAIR principles, in accordance to the guidelines found at INTEGRAPE.

## Methods

### Plant materials and field sampling of grape organs throughout development

Ripening berries belonging to the *Vitis vinifera* cultivars cv. ‘Béquignol Noir’ (some of which presented the variegated berries), and ‘Béquignol blanc’ were sampled from a grapevine germplasm collection (INRA, France) at five weeks after veraison (5WAV) at two consecutive years (for transcriptomic analysis) and every two/three week, starting from the onset of ripening (veraison) for qPCR gene expression analysis and volatile aroma compounds identification and quantification. Around 2 berries/clusters (n=8) belonging to five plants were used for each sample (three biological replicates). Berries were immediately frozen in liquid nitrogen, peeled after slight thawing, and deseeded in liquid nitrogen, to skin and pulp for later analysis. The samples were ground into powder in liquid nitrogen using a ball grinder MM200 (Retsch, Haan, Germany), and stored at –80 °C for later analysis. Grape berries were also harvested from cv. ‘Gewürztraminer’ (at 40, 53, 67, 84, 101, and 116 days after anthesis, DAA) and cv. ‘Viognier’ (32, 45, 73, 92, and 105 DAA) vineyards located in Oliver (BC, Canada). Forty berries per sample were collected for terpene and transcript analyses; berries were cut off from the cluster at the pedicel level, snap frozen with liquid nitrogen, and stored at −80°C. Open flower samples were collected from two-year old vines grown in the UBC Horticulture Greenhouse. At each developmental stage three biological replicates per cultivar were considered. All berries were immediately peeled and deseeded before sample storage All organs and tissues were frozen in liquid nitrogen and stored at −80 °C until required for HPLC analysis or RNA extraction.

### Microscopy

Sections of ripening berries (at 5 weeks after veraison, WAV) were performed with a vibrating-blade microtome (Microm HM 650V). After cutting the fruit in two at the equatorial zone with a scalpel, the half berry obtained was fixed on a metal, placed in the center of the tank microtome. Cross-sections from 60μm to 100μm were cut in water, and then mounted between a blade and a slip with a drop of distilled water. Sections were observed at 10X and 20X with a Zeiss Axiophot microscope and digitalized pictures were obtained with a spot camera (Diagnostic Instruments).

### Phenylpropanoid extractions and HPLC methods

Aliquots of 200 mg of berry skin powder were freeze-dried for 72 h and the dried powders (∼50 mg) were extracted in 1.0 mL methanol containing 0.1% HCL (v/v). Extracts were filtered through a 0.45 μm polypropylene syringe filter (Pall Gelman Corp., Ann Harbor, MI, USA) for HPLC analysis. Each individual sample was analyzed by HPLC as described in (47). Anthocyanin and flavonol quantification were performed as described in (76, 77).

### Molecular marker analysis

Different organs and tissues representing pure L2 (i.e. root and berry pulp) and mixed L1 plus L2 (i.e. leaves and berry skin) cell ontologies were sampled from cv. ‘Béquignol’ somatic variants, including skin samples from variegated berries. Genomic DNA was extracted using a CTAB based extraction procedure slightly modified as described previously (78). The haplotype structure was analyzed using nine different microsatellite markers across distal arm chromosome 2 and analysis of VvMybA1 and VVMybA2 alleles. VvMybA1 gene polymorphisms were investigated using the primers a and d3, and PCR amplifications were performed as reported (79) and F2 and R1 primers described previously (80). PCR fragments were separated by electrophoresis in 1.5% agarose gel in TBE buffer, stained with ethidium bromide and photographed under UV light. For VvMybA2 gene, one-point mutation (SNP) related to berry color, VvMybA2R44 (20), was investigated as described (81) and analyzed by capillary electrophoresis (ABI PRISM 310 Genetic Analyzer, PE Applied Biosystems, California, USA) and data analysis was performed by Peak Scanner Software 2 version 2.0. Microsatellites used were described in different publications: (VvNTM1, VvNTM3, VvNTM4, VvNTM5 and VvNTM6) (82), (SC8_0146_010; SC8_0146_010) Adam-Blondon personal communication, (VMC7G3) (83), (VVIU20) (84). Amplifications were developed separately for each one and size analyses by capillarity electrophoresis were developed in SECUGEN S.L (ABI PRISM 310 Genetic Analyzer. PE Applied Biosystems, California, USA). The analysis of all capillarity electrophoresis data was performed by Peak Scanner Software 2 version 2.0.

### Re-analysis of RNA-seq datasets

Raw sequencing transcriptome datasets were processed using Trimmomatic and filtered reads were aligned to the aligned to the 12X grapevine reference genome using HISAT2 as previously described in (85). Expression of grapevine MYB24 splice variant (MYB24.1 and MYB24.2) were estimated from HISAT2-aligned BAM outputs using Stringtie with default settings. Transcript abundance is expressed as TPM (Transcripts Per Kilobase Million).

### Transcriptomic exploration of variegation

Microarray were produced using the *Vitis vinifera* Array-Ready Oligo Set™ version 1 (Operon Biotechnologies, Germany) as described (86). Microarrays were scanned using a GenePix 4000B fluorescence reader using GenePix Pro version 4 image acquisition software (Axon Instruments, Canada). Spot quantification and quality control was done using Maia version 2.75 (87). Bad quality spots and those with intensities above 50000 were filtered before further analyses. Data analyses were performed using the R/Bioconductor (88) package limma (89). Background correction was done using the normexp method (90). Array normalization was carried out using the limma function ‘normalizeWithinArrays’ (91) and the method ‘printtiploess’. P-Value adjustment was done using the Benjamin-Hochberg method. Genes with expression ratio above 1.6 and adjusted *p*-value below 0.1 were considered as differentially expressed. For probe mapping oligonucleotide probes were mapped to the 12X.1 CRIBI V1 annotation of the PN40024 grapevine genome (92) as in Ensemble Genes (93) release 22. Best matches were considered as targets. Genome sequences were annotated using best blast matches against the uniref100 database, release 15.14 (94) using the qualifiers: homologue to (> 50% alignment identity), similar to (> 70%), weakly similar to (<= 70%), complete (> 98% hit coverage) and partial (<= 98% hit coverage). Functional analyses were done using the MapMan and GSEA Ontology (95, 96) and grapevine mappings made for the used microarray (97). Significantly affected categories were identified using a Wilcoxon rank-sum test implemented in R (R Development Core Team, 2014). Most significantly affected categories were illustrated using R/Lattice (98).

### DNA affinity purification sequencing (DAP-seq)

Genomic DNA (gDNA) was purified from young grapevine leaves of cv. ‘Cabernet Sauvignon’ CS08 by crude nuclei isolation (with PVP40) and 20% Sarkosyl-chloroflorm extraction (99). Genomic DNA library and DAP-seq were performed following published protocol (38). Briefly, the gDNA sample was sonicated into 200 bp fragments on a Covaris Focus-ultrasonicator instrument, which underwent end repair, A-tailing and adapter ligation to attach Illumina-compatible sequencing adapters to the DAP-seq library. We verified successful adapter ligation by qPCR of the DAP-seq library with primer sequences that anneal to the adapter, as well as gel electrophoresis of the sonicated genomic DNA and the DAP-seq library (38). MYB24 was amplified from cv. ‘Cabernet Sauvignon’ and cloned into the pIX-HALO expression plasmid (TAIR Vector:6530264275) to create an expression vector that contained a HALO-tag at the N-terminal. Clones were verified by digestion with restriction enzyme Xho. The MYB24-expression vector was used in a coupled transcription/translation system (Promega) to produce MYB24 protein with the HaloTag-fusion. Expression of MYB24 was confirmed by western blot using an anti-HaloTag antibody (Promega). The protein expression reaction was mixed with HaloTag-ligand conjugated magnetic beads (Promega) to pull down the HaloTag-fused TF. The pulled down TFs was then be mixed with the DAP-seq library for TF-DNA binding to occur. 400ng library was used in each DAP-seq reaction. The bound DNA were then eluted and PCR amplified to generate sequencing libraries that were sequenced on an Illumina NextSeq 500 (sequencing of libraries was set at 30 million and 1×75bp single-end reads). As a negative control, we performed DAP-seq experiment with pIX-HALO expression vector without any ORF insert, accounting for possible non-specific DNA binding in the reaction mixture, as well as copy number variations at specific genomic loci. Sequencing reads were mapped to the grapevine reference genome assembly available at https://cantulab.github.io/data.html. Enrichment of binding events was first evaluated by plotting genome coverage compared to negative control (38). To identify “peaks”, enriched regions relative to control that correspond to binding events, the GEM peak caller was used (100), which performs simultaneous peak calling and motif refinement. *De novo* motif discovery was performed using 200 bp sequences centered at GEM-identified binding events for the 600 most enriched peaks (101). MYB24 target genes were identified by associating the peak regions to TSS of the closet genes.

### Gene co-expression networks generation

Two distinct aggregate gene co-expression networks –condition independent and condition dependent GCNs– were constructed from RNA-Seq datasets downloaded from the SRA database. The condition dependent network only contained datasets extracted from berry tissue from *Vitis vinifera*, while condition independent contained all *Vitis vinifera* available datasets. Networks were constructed as described in (25). Briefly, A total of 131 and 67 SRA studies were obtained, encompassing 2,766 and 1,615 runs from condition-independent and condition-dependent (berry) samples, respectively (Supplementary Table S4A and B). Each SRA study was individually analysed to build a highest reciprocal rank (HRR) matrix (102). To construct the aggregate whole genome co-expression network, the frequency of co-expression interaction(s) across individual HRR matrices was used as edge weights, and after ranking in descending order, the top 420 frequency values for each gene were chosen to build the final aggregate networks. The list of top 420 most highly co-expressed genes (1% of all VCost.v3 gene models) was used to generate individual gene-centred co-expression networks (GCNs). The individual TPSs and MYB24 GCNs extracted from the condition-dependent and -independent networks were analyzed to generate Figure S11.

### Overlap of DAP-seq and transcriptomic datasets

DAP-Seq peaks were converted from cv. ‘Cabernet Sauvignon’ to their cv. ‘Pinot Noir’ corresponding IDs after filtering a BLAST output between both genome annotations (98% identity and 98% coverage thresholds). *TPS* genes were manually curated and assigned to its cultivar counterpart based in BLAST and global alignments. Berry skin transcriptomic datasets from UV-B-irradiated (RMA-normalized) (16) was analyzed using PN40024 12X.2 Assembly V1 annotation, drought stressed (raw data downloaded from PRJNA313234 BioProject at SRA) (17) and ‘light versus shade’ treatments (raw data downloaded from PRJNA661034 at SRA) (43) were reanalyzed using PN40024 12X.2 Assembly VCost.v3 annotation. The RMA-normalized microarray data, containing two developmental stages (23° and 26° Brix degrees) and two UV-B conditions, was compared using a one-factor ANOVA analysis (limma package in R). Genes with (adj pvalue < 0.05 for RNA-seq data, < 0.1 for microarray data, and logFC > 0.53) in at least one berry density were considered as differentially expressed (DE). The Illumina raw sequence reads were trimmed using fastp version 2.0 (the minimum PHRED score accepted for trimming was 20, and reads shorter than 40 bp were discarded). Reads were aligned against the reference PN40024 12X.2 genome assembly using the HISAT2 v2.1.0 aligner with default parameters. Aligned reads were counted with Feature Counts, using the Vcost annotation mapped in 12X.2. Finally, drought and ‘light vs shade’ DE genes (adj pvalue < 0.05 and logFC > 0.53) were extracted using the limma R package. The overlap between UV-B, drought, and ‘light vs shade’ DE genes, the MYB24-GCN and the DAP-seq peaks was performed in R using the UpSetplot package in R.

### Transient tobacco transfection and dual-luciferase reporter assay

The transactivation of the regulative region of TPS35 was tested by Dual-Luciferase Reporter Assay (Promega). The two promoters (TPS35 and HYH from ‘Cabernet Savignon’, CS) and the coding sequences of three TFs (VviMYB24, AtMYC5, and VviMYC2) were amplified and cloned into the entry vector pENTR/D-TOPO (Invitrogen) and then transferred by site-specific recombination into the reporter and effector vectors (pPGWL7.0 and pK7WG2.0, respectively) by using LR clonase. The resulting constructs and the reference vector expressing Renilla luciferase gene (103) were transferred to A. tumefaciens strain EHA105 by heat shock. Dual-Luciferase assays were performed in Agrobacterium-infiltrated tobacco (*Nicotiana benthamiana*) leaves as described by (104). Plants were grown from seeds in a greenhouse with temperature between 30°C and 21°C, relative humidity of approximately 32-50%, a 15 h/9h light/dark cycle, and kept in low light conditions during the whole experiment (three days). Firefly (LUC) and Renilla (REN) luminescence were detected using a GENios Pro TECAN instrument (University of Verona, Biological Institute).

### Nucleic acid extraction and qPCR gene expression quantitation

Different methods of RNA extraction were used depending on plant materials. Total RNA from cv. ‘Cabernet Sauvignon’ and cv. ‘Béquignol’ berry skins was isolated (105). For cv. ‘Gewürztraminer’ and cv. ‘Viognier’, frozen samples were ground to a fine powder under liquid nitrogen using an analytical mill (A11 Basic, IKA). RNA extraction and determination of RNA quality and quantity were performed as described (30). The RT reactions were performed with two μg of total RNA, reverse transcribed with oligo(dT)15 in a 20-μL reaction mixture using the Moloney murine leukemia virus reverse transcriptase (Promega) or the RevertAid First Strand cDNA Synthesis Kit (Thermo Scientific), according to their manufacturer’s instructions. The qPCR reactions were performed for all genes of interest using specified primers. For cv. ‘Béquignol’ samples, the Brilliant® SYBR® Green QPCR Master Reagent Kit (Stratagene) and the Mx3000P detection system (Stratagene) was used. In the rest of cases, PowerUp SYBR Green Master Mix (Thermo Scientific) was used with an ABI 7500 Real-Time PCR System (Applied Biosystems). Amplification of the Vvi*UBIQUITIN1* gene (VIT_16s0098g01190, 99bp) (106), *VviELONGATION FACTOR1* (*EF1;* VIT_00025941001, 91bp) and *VviACTIN* (VIT_00026580001, 81bp) (107) were used for normalization of relative expression measures. All RT qPCR biological replicates were run in two or three technical replicates within the same plate. PCR conditions, standard quantification curves for each gene and relative gene expression calculations were conducted (30).

### Volatile determination in cv. ‘Gewurztraminer’ and ‘Viognier’

Free (non-glycosylated) VOCs were analysed with some modifications (108). Five g of frozen grape powder were weight out in a 20 ml SPME glass screw cap vial with 1.5 g sodium chloride. Then 4 ml citrate phosphate buffer (pH 5.0) and 100 µl 200 g/L ascorbic acid were added. Finally, 3-Octanol (10ppb final concentration) and d3-Linalool (10ppb final concentration) were added to the sample as internal standards. The free volatile compounds were extracted and analyzed by headspace–solid phase microextraction (SPME) gas chromatography-mass spectrometry (GCMS). A carboxen–divinylbenzene–polydimethylsiloxane SPME fiber was used (No. 57329-U Supelco, Sigma, St Louis, MO, USA). VOC analyses were performed on a gas chromatograph (model 7890A, Agilent, Waldbron, Germany) equipped with a 5975C mass spectrometer detector and GC PAL80 autosampler (Agilent Technologies). The column used was an Agilent J&W Scientific DB-WAX (30m x 0.25mm ID with 0.25um film thickness) with a film thickness of 0.25 mm. Helium was used as the carrier gas at a constant flow of 0.8mL min^-1^. The capped vials were heated at 40 C for 20 min to promote the transference of the compounds from the sample to the headspace. After this step, the SPME fiber was inserted into the sample vial headspace (22mm depth) for a 30 min sample extraction at 40 C and then inserted into the GC injection port at 250 C and kept for 5 min for the desorption. The injection port was lined with a 0.75-mm splitless glass liner. ThePulse splitless injection mode was used. The oven temperature program was as follows: 40 C for 4 min, followed by 3 C /min up to 150 C, 25 C /min to 230 C, 230 C for 10 min. The mass spectrometer was operated in the electron impact mode at 70 eV, scanning from *m*/*z* 33–500. The ion source temperature was 230 C and the quadrupole temperature was 150 C. Data processing was carried out by MSD Chemstation (E.02.02.1431, Agilent Technologies Inc.). Only VOC peaks with signal-to-noise ratios greater than 10:1 were considered for data quantification. Identification was achieved by comparing the liner retention index (LRI) and the mass spectra present in Wiley09Nist08 Database (Agilent Technologies Inc.). Compounds were semi-quantified using d3-linalool as internal standard.

### GC-MS analysis of volatiles in cv. ‘Béquignol’ samples

Samples were prepared and solid-phase extracted (SPE). Three biological replicates of red and white skins of variegated berries were used. Skins were grounded under liquid nitrogen and suspended in 4 mL of a sodium sulphite solution at (Na_2_SO_3_, 10 g/l). The supernatant obtained after a 15 min centrifugation (11400 x g at 4 °C) was passed through a glass fibre prefilter and a glass filter. Twenty microliters of a 1 g/L 3-octanol solution were added as external standard to allow for the quantification of the main terpenols. The filtrate was passed through a 1 g phase C18 silica-bonded non-polar column (HF Mega Bond Bond Elut C18, Agilent, 5301 Stevens Creek Blvd, Santa Clara, CA 95051, United States), previously rinsed with 6 mL of methanol and 2×6 mL of ultrapure water, at a rate of approximately 1 drop/s. Total fractions of free and bound monoterpenols were eluted with 4 mL of absolute ethanol. This total terpenol extract was diluted in 40 mL of citrate-phosphate buffer (pH 4.5) and the resulting solution was incubated with 50 mg of Rapidase® AR 2000 glycolytic enzyme (DSM Food Specialties Beverage Ingredients P.O. Box 1, 2600 MA Delft, The Netherlands) overnight at 37.5 °C to release the glycosidically bound terpenols. The released terpenols were separated by SPE again on a C18 column and eluted with 4 mL of dichloromethane after rinsing with water. Extracts were dried in Pasteur pipettes filled with 0.5 g of anhydrous sodium sulphite and concentrated to 500 µl under a gentle nitrogen flux. Twenty microliters of a 1 g/L m-cresol solution were added to each sample as internal control. Samples were stored at –20 °C prior to gas chromatography analysis. For GC-MS non-targeted analysis, chromatogram files were deconvoluted and converted to ELU format using the AMDIS Mass Spectrometry software (http://www.amdis.net), with the resolution and sensitivity set to medium. Chromatograms were then aligned and integrated using Spectconnect (http://spectconnect.mit.edu). All metabolites found in the blank run or believed to have originated from the column bleeding were removed from analysis at this time. After removal of contaminant metabolites (volatiles), the IS matrix from Spectconnect was normalized according to the sample weight and 3-octanol surface (external standard). Volatiles were identified by comparison to the NIST 14 standard reference database. Identities of metabolites of interest were then confirmed by authentic standards when available. Terpenols were quantified through GC-MS targeted analysis. Extracts were analyzed by GC-MS using an Agilent 6890 gas chromatograph equipped with a Gerstel MP2 autosampler and an Agilent 5973N mass spectrometer for peak detection and compound identification. The GC was fitted with a DB-Wax column (30m×0.32mm i.d., 0.5 µm film thickness, Agilent J&W, Agilent Technologies France, 3, Avenue du Canada, 91978 les Ulis France). Helium was used as carrier gas with a column flow rate of 1.5 mL/min. The GC oven temperature was programmed from 45°C to 235° C, at 2.7°C/min (hold 10 min). The injector was set to 230°C and used in pulsed splitless mode (15 psi for 0.50 min). The MS transfer line and ion source temperatures were set at 270 °C and 230 °C, respectively. The MS was operated in EI mode and positive ions at 70 eV were recorded with a scan range from m/z 30 to m/z 400. Agilent MSD ChemStation software (G1701DA, Rev D.03.00) was used for instrument control and data acquisition. Total amounts of 3-octanol, m-cresol, alpha-terpineol, citronellol, linalool, nerol and geraniol were determined using linear calibration curves built with standard, GC quality grade, molecules over a concentration range from 0 to 100 ng/µL. Final concentrations were expressed in microgram/l.

### Sunlight/UV-B experiments

Three different sunlight or UV-B radiation experiments were conducted in cv. ‘Cabernet Sauvignon’ commercial plants (either treated at field or uprooted and transferred to a greenhouse). Experiment 1; low fluence UV-B exposure treatment applied to cluster from nine-year-old potted vines in a UV-free greenhouse during 2011-2012 and 2012-2013 growing seasons (n=3), respectively (45). Experiment 2: A UV-B filtering radiation treatment was applied in a commercial vineyard as described previously (46) during 2011-2012 growing season (n=4). The filtering treatment consisted in blocking solar UV-B radiation by installing a 100 µm clear polyester film at the position of grape clusters. Experiment 3: Sunlight reduction treatments were conducted at field as described previously (11) and consisted in full shading of fruits by the plant’s own canopy (0% exposure) and full sunlight exposure from veraison (ripening onset) onwards, generated by displacement of leaves around the cluster region (100% exposure, n=3). Berry skins were frozen in liquid nitrogen and stored at −80°C until RNA was extracted.

### Light shading experiments in cv. ‘Gamay’ and cv. ‘Gamay Fréaux’

Light exclusion was conducted from vines of 23-years old, spur pruned, with a density of 1.6 m between rows and 1 m between plants in a germplasm collection vineyard. Eighteen vines were chosen to form three blocks with three vines each and two similar clusters from two adjacent shoots of each vine were tagged. Opaque boxes were applied to one of the two tagged clusters of each vine from 2 weeks before veraison until maturity for light-exclusion treatment, and the other clusters were exposed under natural light conditions as the control as described previously (47). Light-exposed or shaded clusters of both cultivars were sampled weekly from one week after treatment until maturity. Three berries from each cluster and three clusters from three vines per block were sampled for each treatment at each sampling date. Berries were weighed, deseeded, separated into skin and pulp, and the skin and pulp were immediately frozen in liquid nitrogen. The samples were ground into powder in liquid nitrogen using a ball grinder MM200 (Retsch, Haan, Germany), and stored at –80 °C for later analysis.

### Transactivation Assay in yeast

The coding sequence of MYB24 was cloned into the pDEST32 vector by LR recombination to obtain the *Gal4DBD:VviMYB24* construction. The fusion protein vector was transformed into the *Saccharomyces cerevisiae* strains SFY526 (109) and NLY2:SS-38 (110). Transformants were selected on SD-glucose medium supplemented with -Leu drop-out solution (BD Biosciences). Confirmation of interaction was performed by targeted transformation of the specific constructs using the small-scale yeast transformation protocol as described in the yeast protocol handbook (Clontech). Transformants were grown in YPDA media to OD600 de 0,5 – 0,8. Tubes with1,5 mL of culture media were centrifuged at 1400 rpm by 30 sec and the supernant was discarded. The pellet was suspended by pipetting in 300µL of Z-buffer. Then the solution was frozen in liquid nitrogen and thawed in 37°C by 30 sec and repeated 3 times. Finally, transcriptional activity was measured using ortho-nitrophenyl-β-galactoside (ONPG) as substrate for β-galactosidase enzyme released from SFY526 and NLY2:SS-38 yeast strains transformed with *Gal4DBD:VviMYB24*. When hydrolyzed, ONPG generates a galactose molecule and an o-nitrophenol molecule, with the latter being quantified by spectrophotometer at a wavelength of 410nm.

### Confocal microscopy

*Agrobacterium tume*faciens strain GV3101, previously transformed with the vector pK7FWG2 (35S:VviMYB24-eGFP), was grown in 10 mL of liquid LB supplemented with rifampicin 50 ng / L, gentamicin 25 ng / L and spectinomycin 50 ng / L. Bacteria isolated from the selection medium were resuspended in 4 mL of 10mM MgCl2 to be infiltrated. Tobacco leaves were infiltrated through the abaxial face with a needleless syringe. The plants were left in the greenhouse for 3 days. Thirty minutes prior to visualization, 1 cm^2^ leaf sections were incubated in a 20 mg / mL solution of 4’, 6-diamino-2-phenylindole (DAPI). Fluorescence was visualized with a Nikon Confocal Eclipse C2si microscope. A 405 nm laser was used to excite DAPI and a filter cube with emission at 445/35 (427 - 462nm) was used for its detection. For GFP, a 488 nm laser and a filter cube with emission at 525/50 (500 - 550nm) and 600/50 (575 - 625nm) were used.

### Bimolecular fluorescence complementation (BiFC) assay

The BiFC assays were performed in Agrobacterium-infiltrated tobacco (Nicotiana benthamiana) leaves as described (111). The four TFs (VviMYB24, AtMYC5, VviMYC2, and VviMYC1) were amplified and cloned to two binary vectors (35S::NtGFP and 35S::CtGFP). The recombinant constructs were transferred into A. tumefaciens strain C58 by heat shock. A single agrobacterium transformant was used to inoculate overnight with antibiotics. The final culture was centrifuged at room temperature with low speeds. The pellets were suspended with MMA solution (0.1 M PH=5.6 MES, 1 M MgCl2, 0.1 M Acetosyringone). According to the experimental combination, the suspension was diluted and mixed, keeping the OD600 as the 0.2 for each construct. The mixture was infiltrated on the abaxial side of leaves. Plants were grown at normal condition (16 h light at 25 °C and 8 h dark at 22 °C). After 48h, infiltrated samples were collected without leaves nerves. Samples were as observed by a Zeiss LSM780 AxioObserver confocal microscope at 40X. GFP (Exc 488nm, Emission 490-544nm) and Chlorophyll (Exc 488nm, Emission 680-760 nm).

### *MYB24*/*HYH* correlation with metabolite composition in berry skins of drought stressed plants

Transcriptomic data (17, 44) was remapped to the PN40024 12X.2 assembly and VCost.v3 annotation. Genes with low expressions were filtered according to the method described in (112), and raw counts were normalized to FPKM (Fragments Per Kilobase Million) values. Applying the Weighted Gene Co-Expression Network Analysis (*WGCNA*) R package (113), the genes were grouped in different clusters. The parameters used were a blockwiseModules with a soft-threshold power value of 30 (fitting a scale free topology network), a deepSplit of 4 and a mergeCutHeight of 0.2, obtaining a total of 28 modules. The correlation between gene expressions or the modules eigengenes (1st principal component) and the metabolite quantification was visualized using the pheatmap (https://CRAN.R-project.org/package=pheatmap) R package, performing a hierarchical clustering using the complete linkage method with Euclidean distance.

