## Supplemental Figures for "The grape MYB24 mediates the coordination of light-induced terpene and flavonol accumulation in response to berry anthocyanin sunscreen depletion"

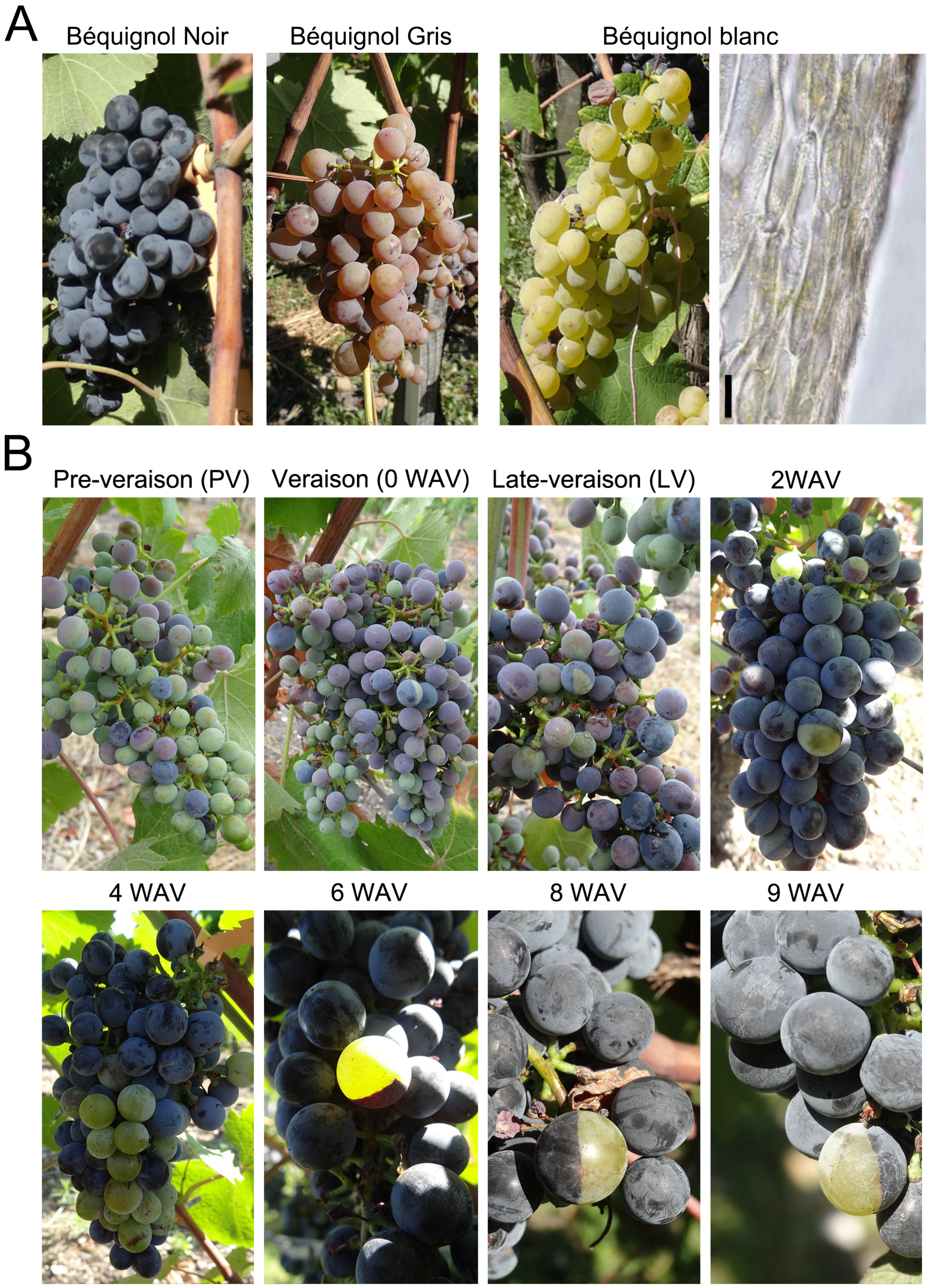


Figure S1: (A) Béquignol color somatic variants. Light field microscopy images of cv. 'B. Blanc' berry skins at full maturation (8WAV) are also shown. Bar scale: 5 µm. (B) Ripening progression at field in the variegated cv. 'Béquignol Noir'. Full maturity is at 8WAV. At 9WAV, berries showed dehydration symptoms characteristic of over-ripening.

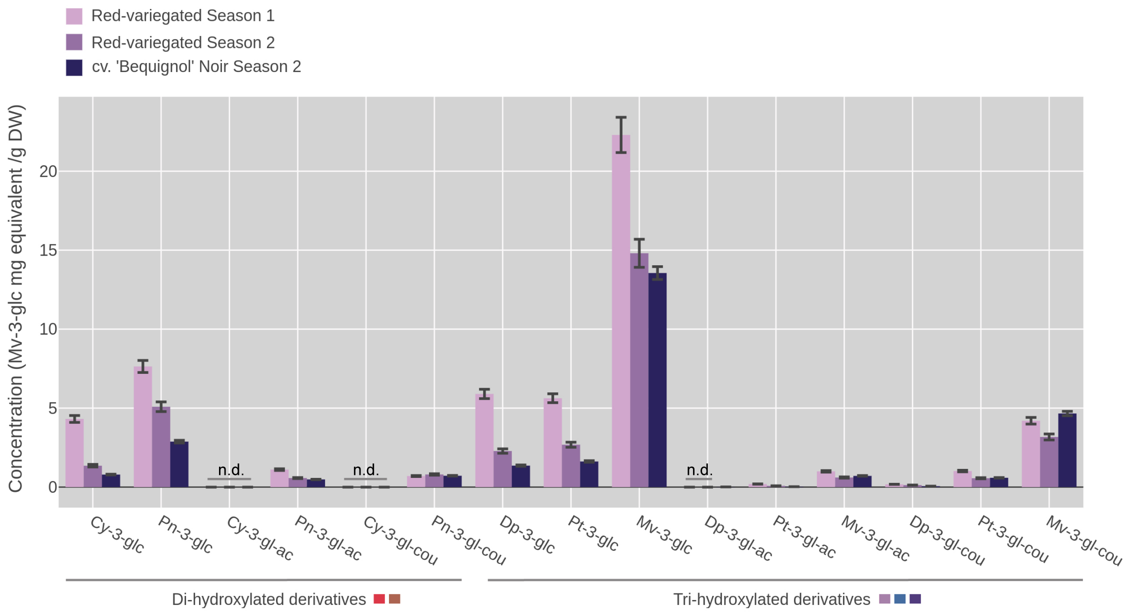


Figure S2: Anthocyanin composition in variegated and non-variegated cv. 'Béquignol Noir' berry skins at 5WAV. High performance liquid chromatography (HPLC) quantifications are expressed as mg/g of dry weight (DW). Standard error bars were calculated from biological replicates. Anthocyanin derivatives are ordered according to their 3'(di)- or 3'-5'(tri)-hydroxylation pattern, namely cyanidin (Cy), peonidin (Pn), delphinidin (Dp), Petunidin (Pt) and Malvidin (Mv).

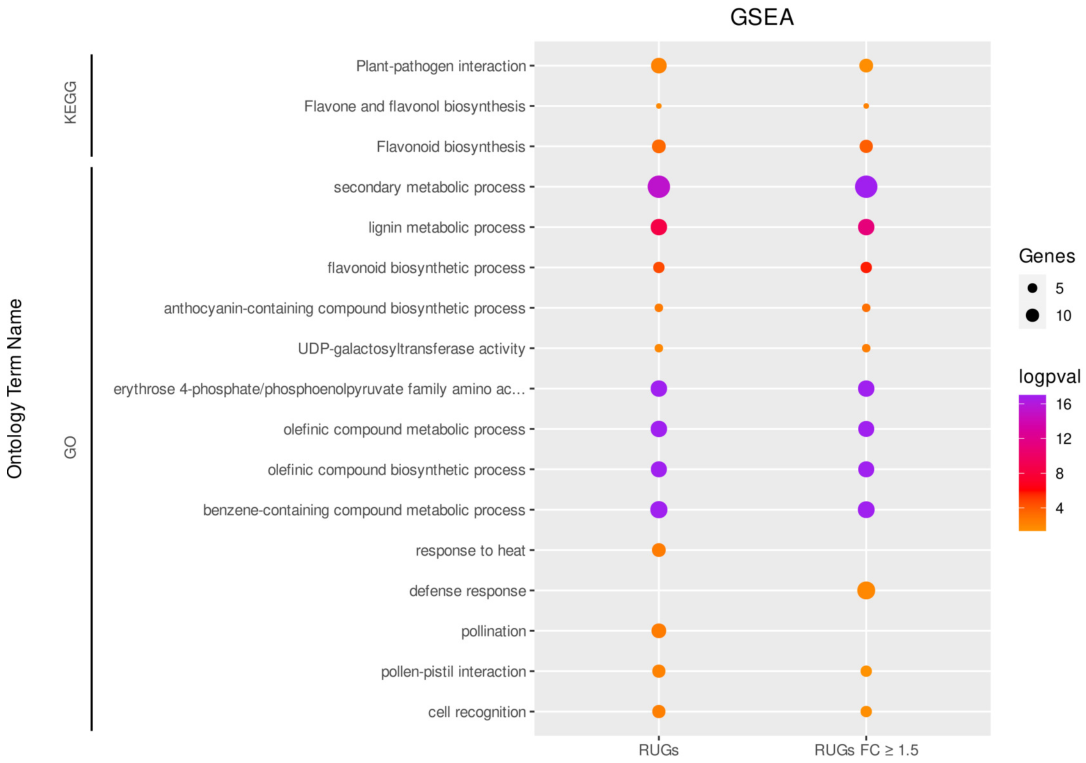


Figure S3: A selection of significantly enriched terms in red-skin up-regulated genes (RUGs). RUGs are in the Supplementary Table S1D and E ($foldchange \geqslant 1.5$) and the complete list of gene set enrichment analysis is in the Supplementary Table S1G and H ($foldchange \geqslant 1.5$).

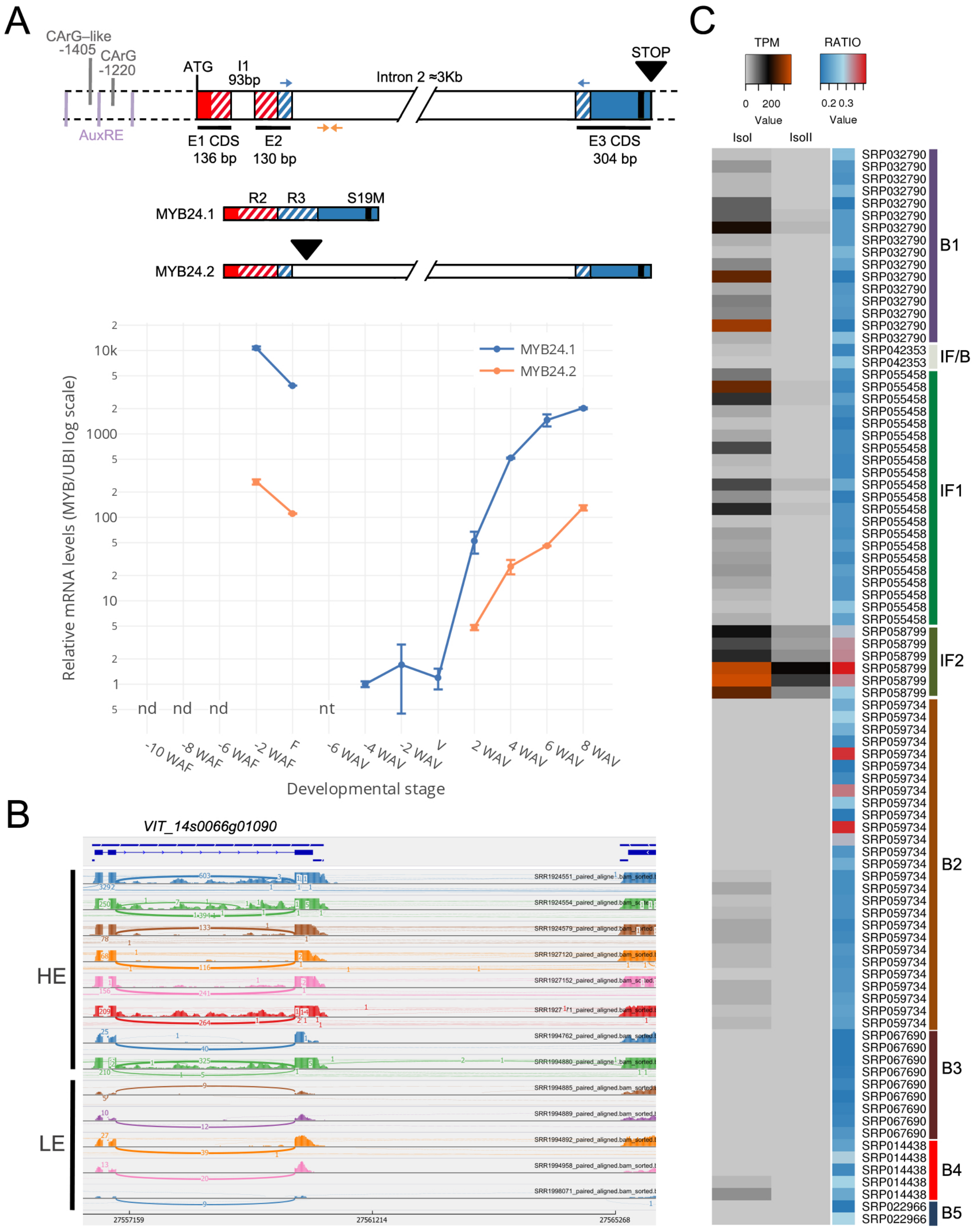


Figure S4: Expression domains of MYB24 splicing variants. (A) Upper panel: Gene structure organization of MYB24. Distances of transcription factor binding sites in the 5' regulatory region are indicated with respect to the start codon (ATG). Arrows indicate primer position for the amplification of isoforms IsoI (MYB24.1) and IsoII (MYB24.2). Bottom panel: expression of MYB24 isoforms in flower and fruit developmental stages of cv. 'Cabernet Sauvignon' grown at field. F: flowering, nd: not detected, nt: not tested. (B) Sashimi plots for a collection of RNA-seq datasets in which MYB24 is highly (HE) or lowly (LE) expressed. (C) Quantification of isoform ratios derived from inflorescence (IF) and berry (B1-5)-specific RNA-seq developmental datasets. TPM: transcripts per million.

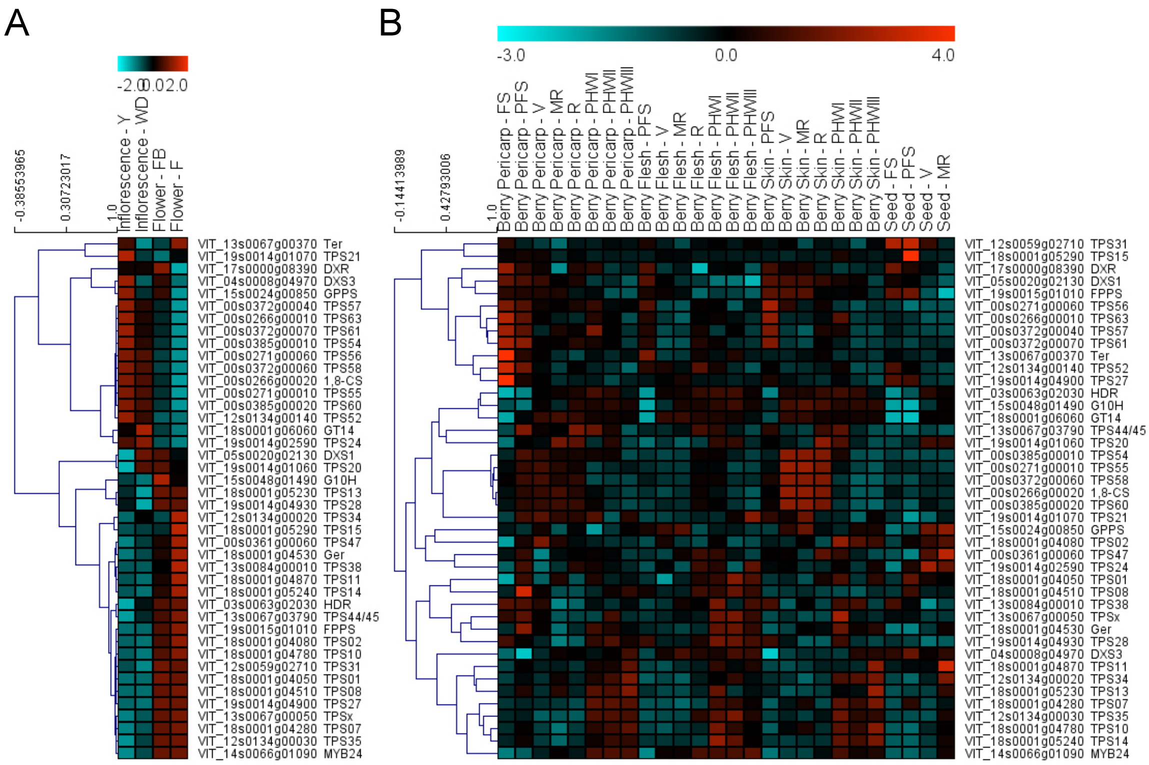


Figure S5: Several grape sesqui- and mono-terpene synthase (TPS) genes are highly co-expressed with MYB24 in late stages of (A) flower and (B) berry development. (A-B) Gene expressions retrieved from the microarray (Nimblegen) cv. 'Corvina' expression atlas, with stages abbreviated as in Fasoli et al. (2012). The gene expression data were calculated as log2, and genes were hierarchically clustered based on average Pearson's distance metric. Red and cyan boxes indicate high and low expression levels, respectively, for each gene.

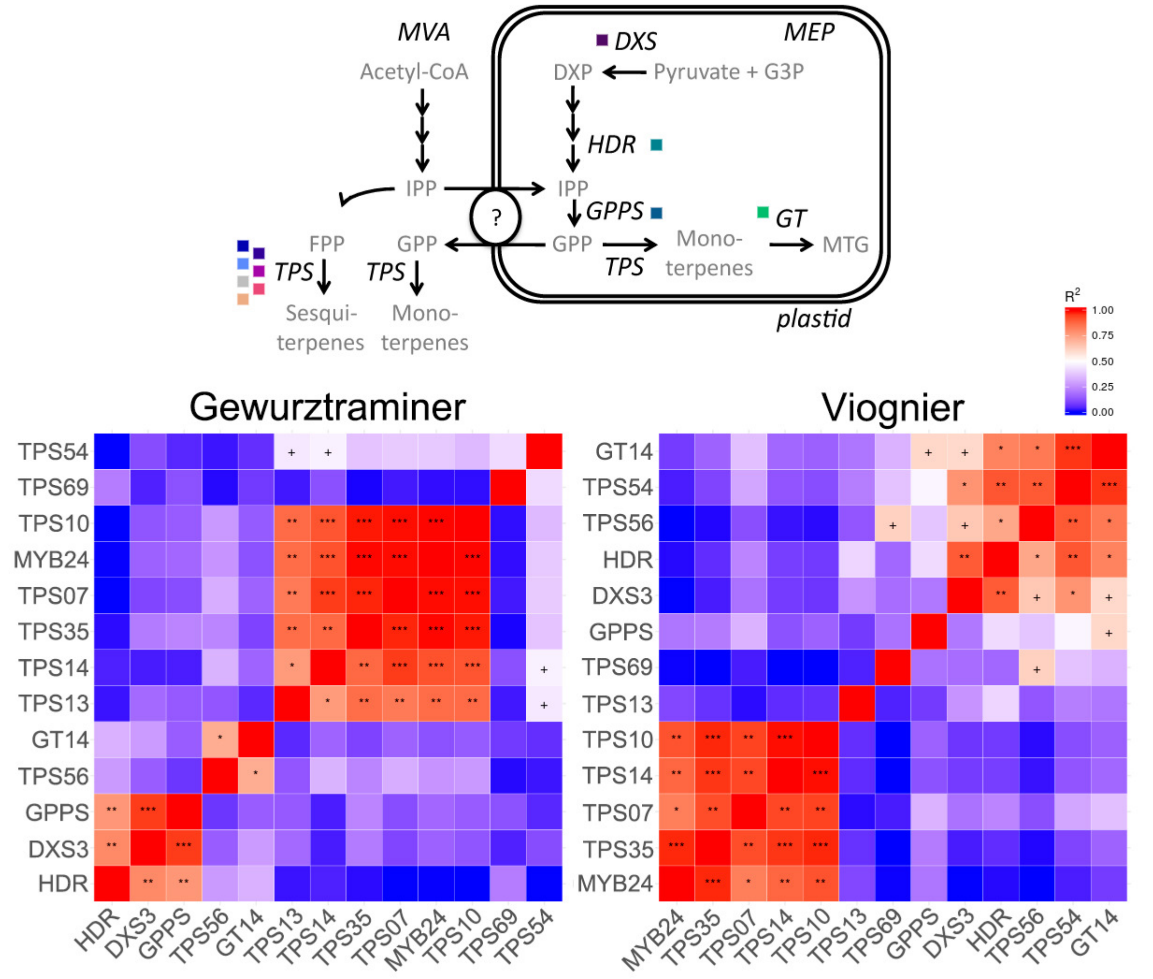


Figure S6: Correlation matrices for MYB24 and isoprenoid/terpenoid gene expressions in flower/berry of the high and low terpene-accumulating 'Gewurztraminer' and "Viognier' cultivars, respectively. Enzymes coded by these genes are referenced to the MEP and MVA pathways (upper panel). (+), (*), (**) and (***) symbols for p<0.1, p<0.05, p<0.01 and p<0.001, respectively.

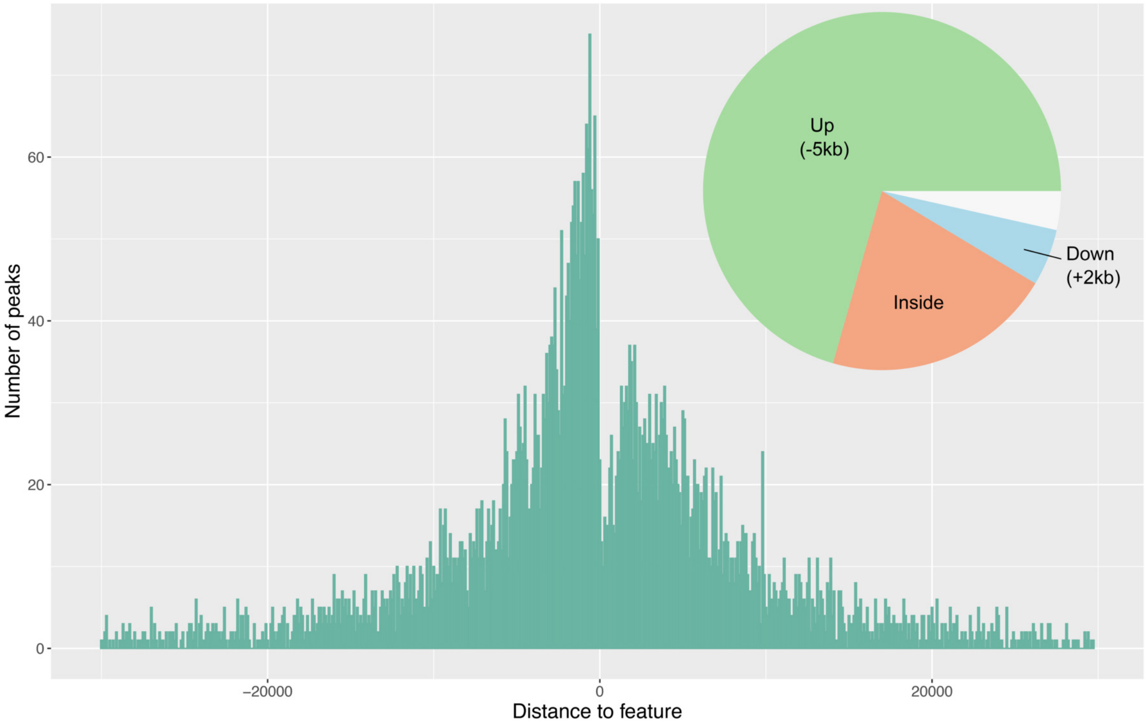


Figure S7: Distribution of MYB24 peaks in Cabernet Sauvignon CS08 v1.0 genome with respect to all transcription start sites (TSSs) of assigned genes. The proportion of binding peaks 5kb upstream of TSSs, inside genes or 2kb downstream of genes are represented within the pie-charts in green, orange and blue, respectively.

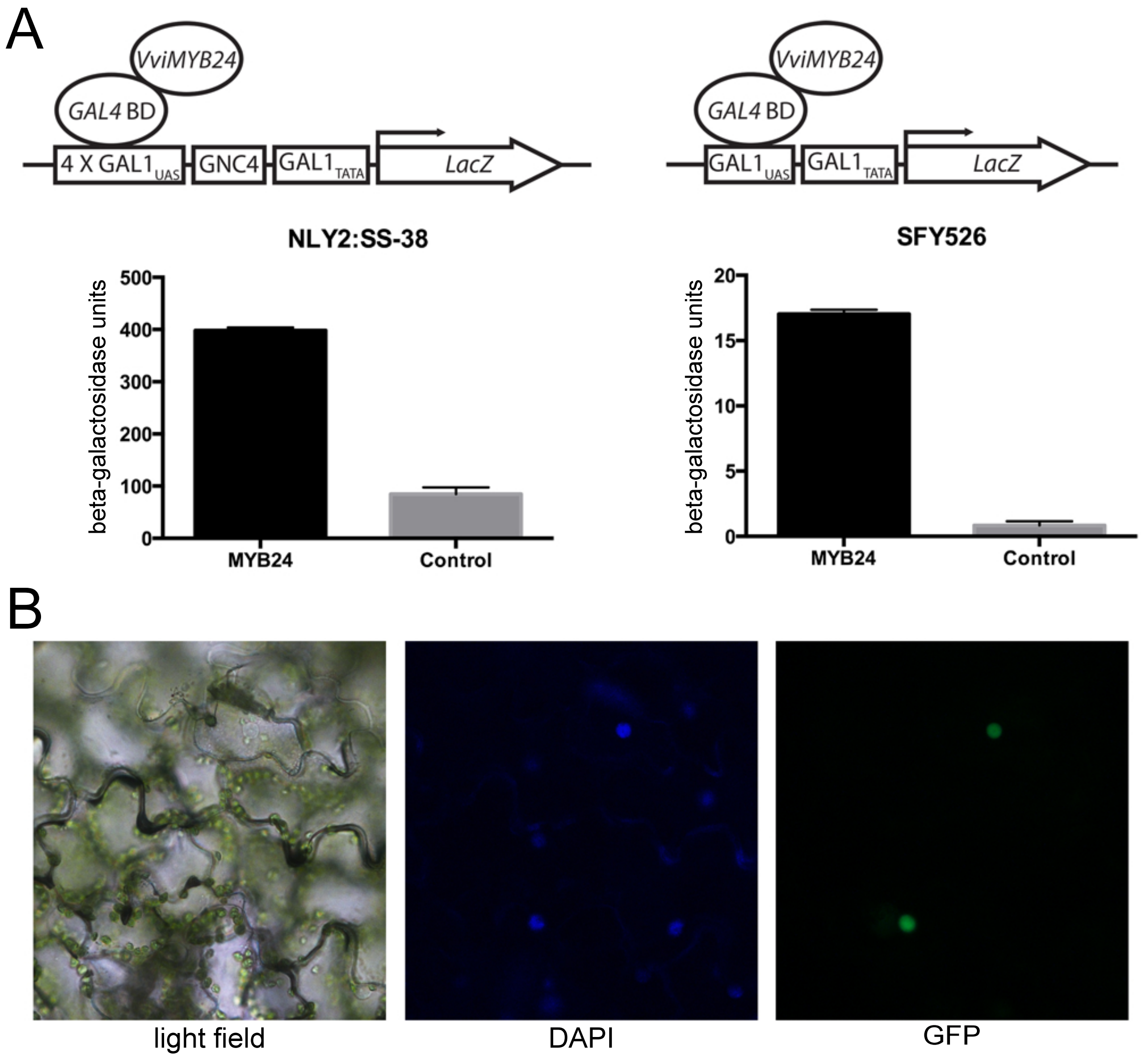


Figure S8: MYB24 is a nuclear-localized transcriptional activator *in vivo*. (A) $\beta$-galactosidase assays of transcriptional activation of GAL4-MYB24 in yeast. The reporter strains used were SFY526 (with an inducible system that has a GAL4BD-binding promoter domain -GAL1UAS- fused to $\beta$-galactosidase LacZ) and NLY2: SS-38 (for testing both repression and activation, with a constitutive expression system of several GAL1UAS binding boxes and two boxes of binding to Gcn4 -GCN4- that allow constitutive expression of LacZ). Enzymatic activity (hydrolysis of 2-Nitrophenyl $\beta$-D-galactopyranoside) of yeast extracts were taken with three biological replicates. (B) Sub-cellular localization analysis of VviMYB24 in Nicotiana tabacum, agroinfiltrated with a 35S:MYB24-eGFP construct (vector pK7FWG2). Three days after agroinfiltration, leaves were incubated with the nuclear marker 4',6-Diamidino-2-phenylindole (DAPI) and later visualized in a confocal fluorescence microscope. Picture is representative of five independent agroinfiltrations.

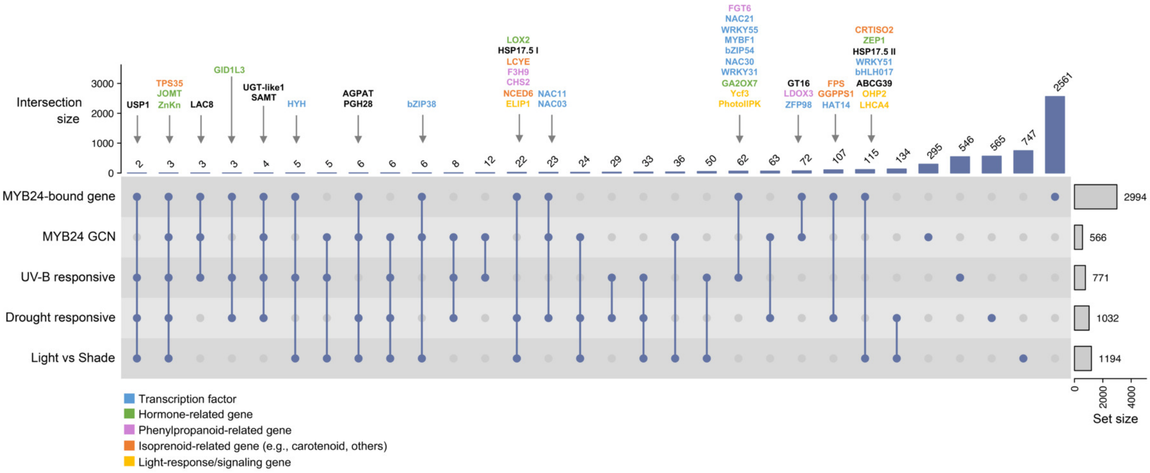


Figure S9: Definition of MYB24 Target genes, resulting from the overlap of DAP-seq data (mapped on the PN40024 12X.2 assembly), MYB24 co-expression data (merge of condition-dependent and -independent networks) and upregulated genes in UV-B radiation, drought stress, and 'light vs shade' transcriptomic studies. A selection of different types of genes are shown for each intersection.

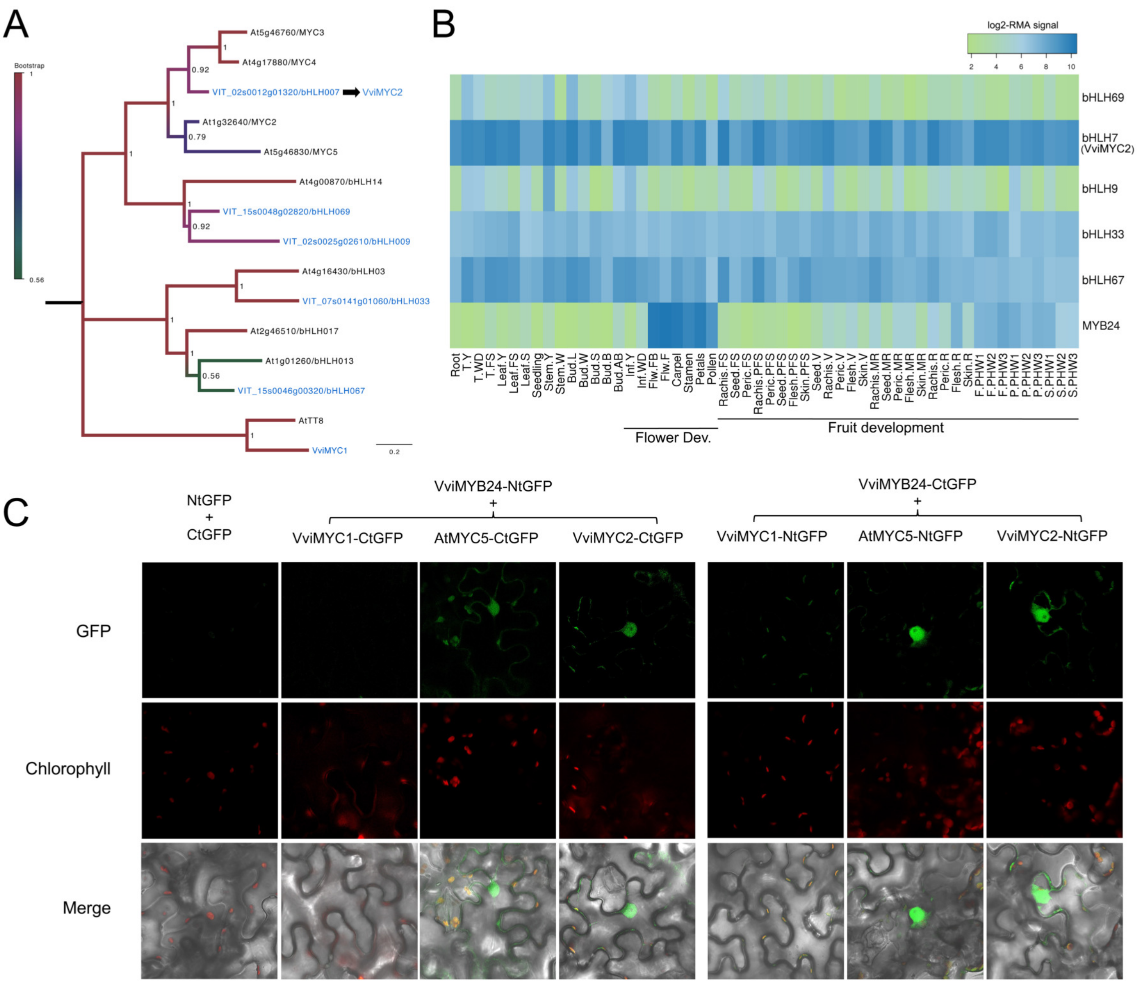


Figure S10: (A) Maximum likelihood tree (2000 bootstraps) with phylogenetic relationships of grape and *Arabidopsis* bHLH proteins from Subgroup 8 (defined in Toledo-Ortiz et al., 2003). Subgroup 7 bHLH proteins involved in flavonoid synthesis were included as outgroup (AtTT8/VviMYC1). (B) Expression behavior of MYB24 and Subgroup 8 bHLHs in the cv. 'Corvina' atlas. (C) Bimolecular fluorescence complementation (BiFC) assay, for testing the interaction of VviMYB24 and bHLHs proteins (AtMYC5 and VviMYC1/2). NtGFP and CtGFP correspond to empty constructs used as negative controls, containing N- (NtGFP) and C- (CtGFP) terminal segments of the green flurescent protein (GFP). VviMYB24 and MYCs were fused to both NtGFP and CtGFP and agroinfiltrated in combination to test interactions. The combination of VviMYB24 and VviMYC1 was used as an expected negative control. The green fluorescence was observed at two days after agroinfiltration in Nicotiana benthaminana leaves. The GFP signals were captured by a Zeiss LSM780 AxioObserver confocal microscope at 40X (Exc 488nm, Emission 490- 544nm).

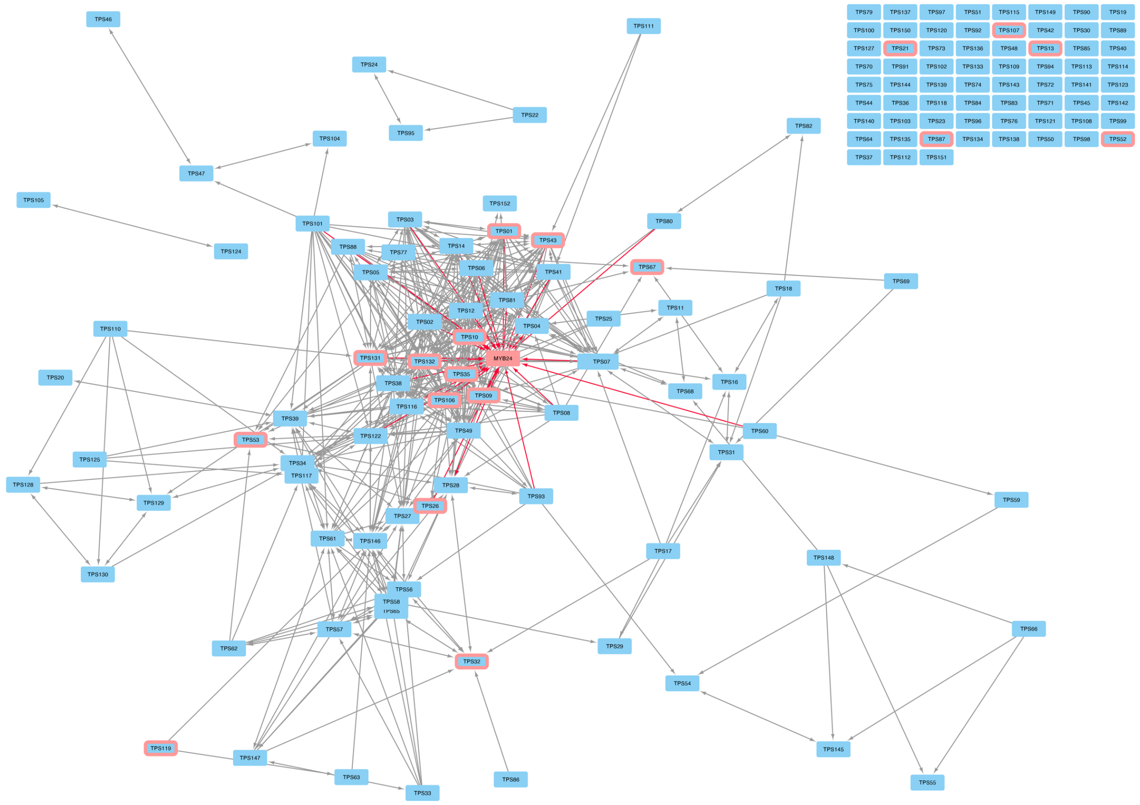

Figure S11: Regulatory and reciprocal co-expression network of MYB24 and the grape terpene synthase family. Co-expression relationships are shown among all genes, when present. Grey arrows refer to gene positive and significant co-expression between all nodes, while red arrows represent the specific co-expression between MYB24 (red node) with TPS genes (blue nodes). Pink node borders indicate MYB24 binding in TPS gene promoter. GCN was generated as in Orduña et al., (2021) from condition-independent and condition-dependent (131 fruit/flower SRA public studies: 2,766 runs) aggregate networks. Network visualization is based on the AggGCN app (https://tinyurl.com/2s3c4z73) developed at the Vitis Visualization platform (Navarro et al., 2021).

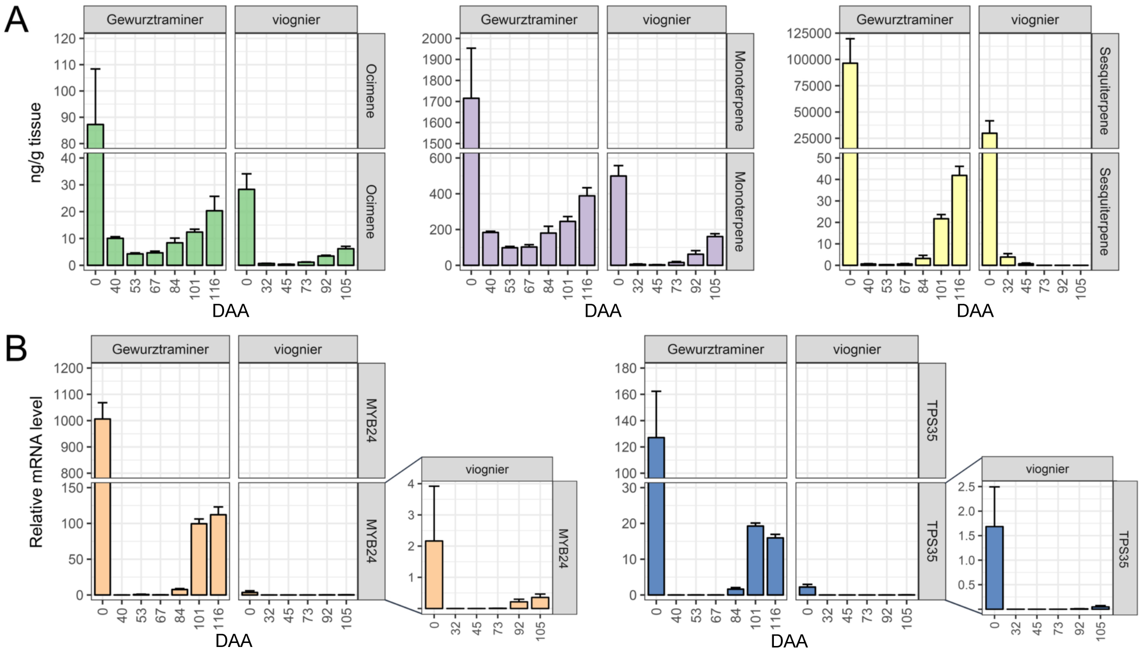


Figure S12: (A) Developmental accumulation of terpenoid volatile compounds in the high terpene-accumulating cultivar cv. 'Gewurztraminer (GW)' and the low terpene-accumulating cv. 'Viognier' (left panel: beta-ocimene; middle panel: monoterpene total content; right panel: sesquiterpene total content). (B) The expression of *MYB24* and *TPS35*. Stages correspond to days after anthesis (DAA). DAA '0' corresponds to blooming, while veraison was 62 and 67 DAA for cv. 'GW' and 'Viognier', respectively. Standard error bars were calculated from biological replicates.

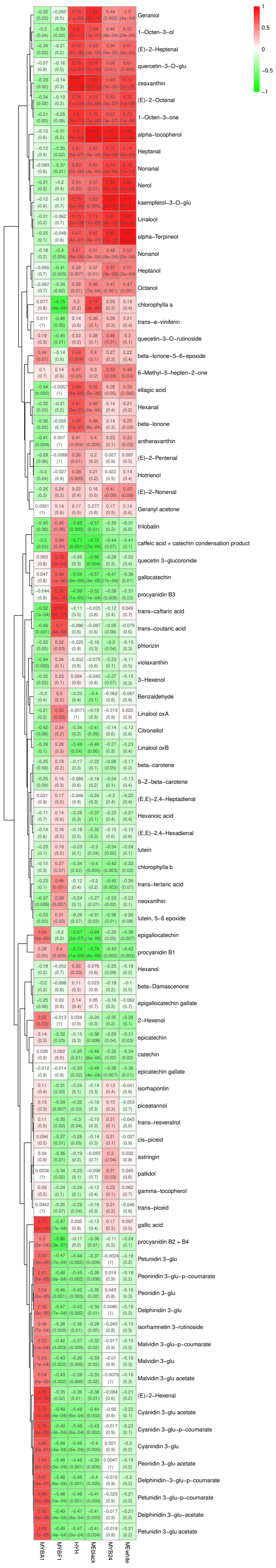


Figure S13: legend (next page)

Figure S13 (previous page): Correlation analysis between specialized metabolites and mRNA abundances deduced from the integration of Savoi et al. (2016) and Savoi et al. (2017) transcriptomics/metabolomics datasets. Rows correspond to metabolites while columns refer to TFs and eigengene of modules (ME) where these TFs are present (ME-white for MYB24 and ME-black for HYH). MEs were obtained by joining expression data from both studies (belonging to white-skin cv. 'Tocai' and red-skin cv. 'Merlot' berry samples) and running a WGCNA clustering analysis. The values in each cell depicts the corresponding Pearson correlation with its significance (p-value) in brackets.

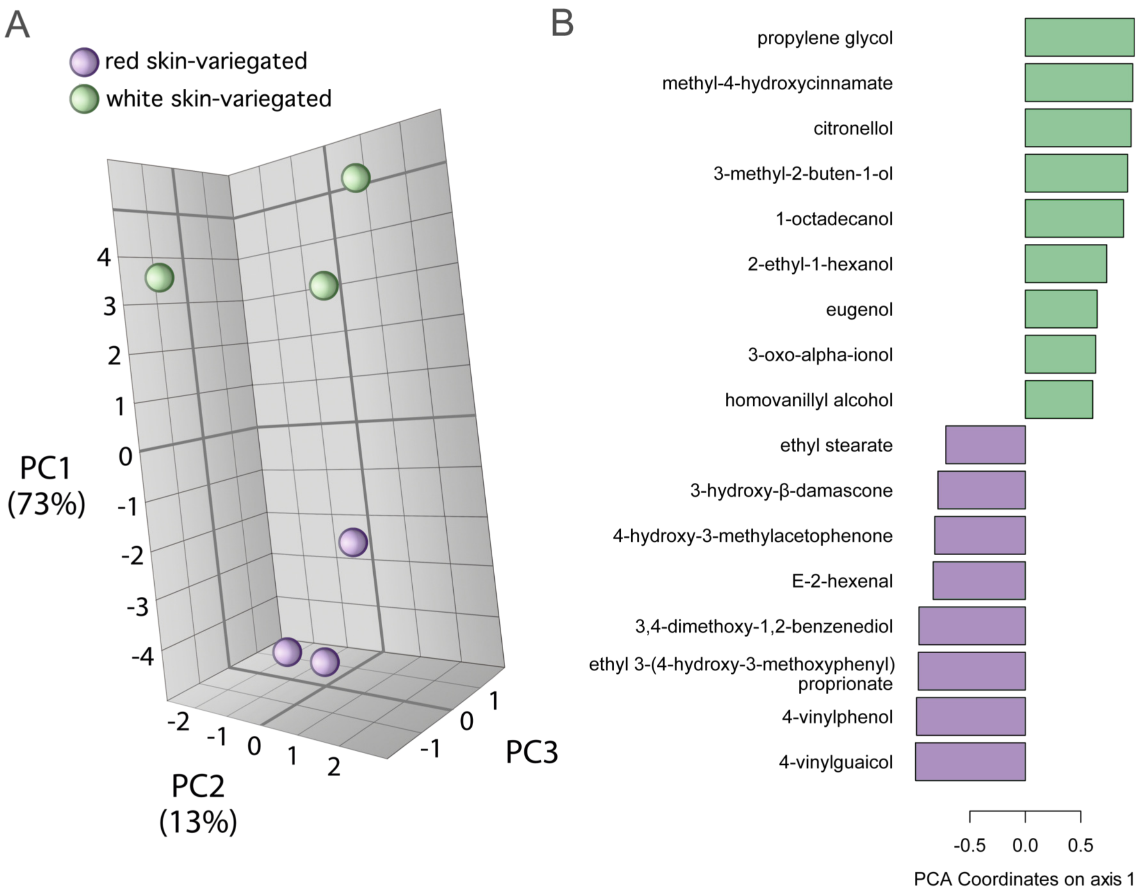


Figure S14: Gas chromatography–mass spectrometry (GC-MS) untargeted volatile compound analysis in red and white skin sections of the variegated cv. 'Béquignol Noir' berry. Berry skin samples were treated with a solid phase extraction (SPE) method. (A) Principal component analysis for berry skin samples collected at 8WAV. (B) Analysis of significantly altered volatile compounds within PC1.

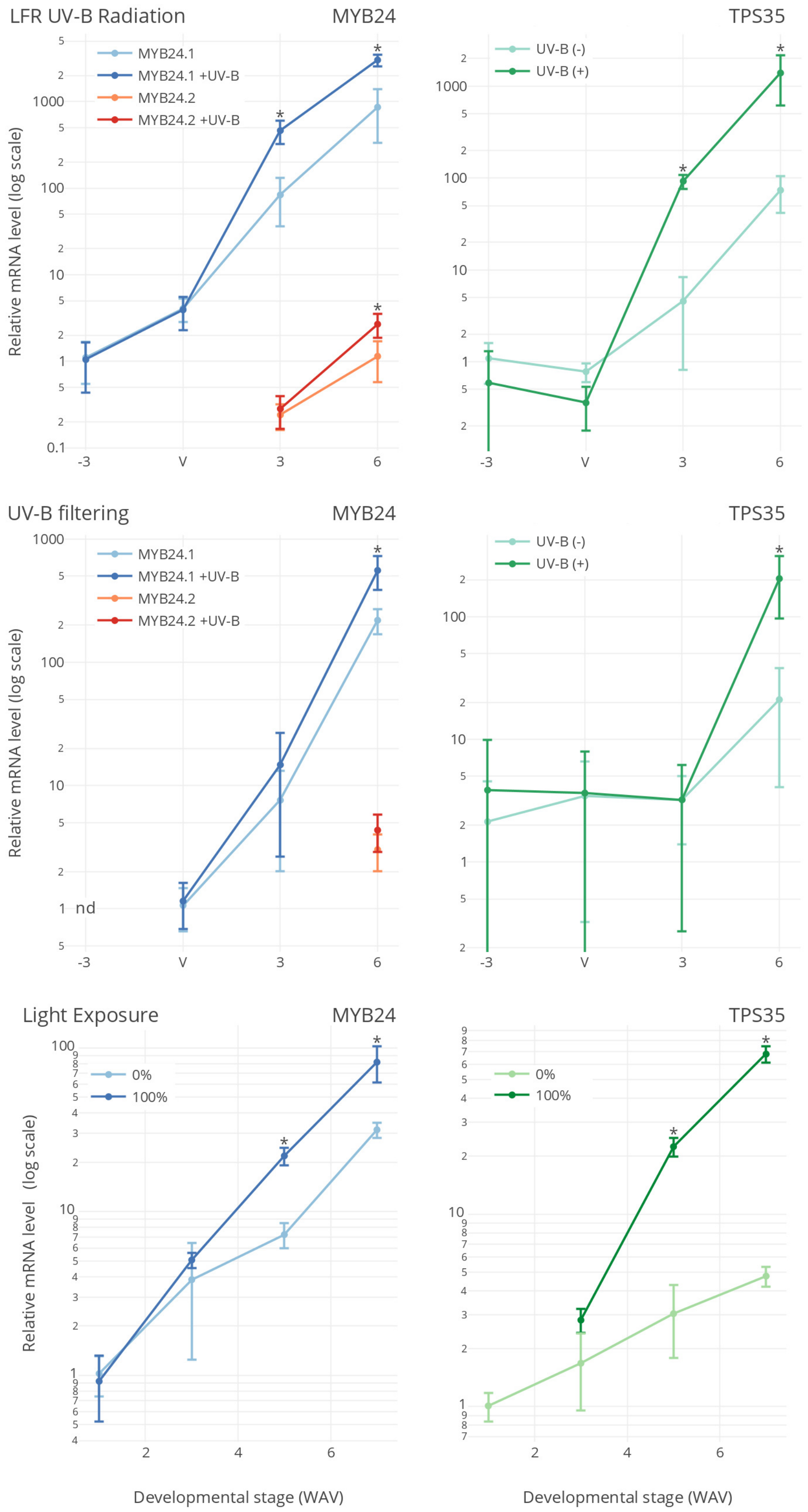


Figure S15: legend (next page)

Figure S15 (previous page): MYB24 and TPS35 expression responses to UV-B irradiation and sunlight/UV-B depletion in glasshouse and field trials. Experiments consisted in i) low fluence rate (LFR) UV-B, directly irradiated on fruits of 9-year-old potted vines, ii) UV-B filtering of fruits at field and iii) fruit sunlight exposure introduced by leaf displacement at field. Sunlight percentages for each treatment refer to the range of time under sunlight exposure: 0% corresponds to full shading of fruits by plant leaves, 100% corresponds to full sunlight exposure from veraison onwards, caused by movement of leaves around the cluster region. The experimental design consisted in four blocks with five plants each (biological replicates n = 4). Three berries per cluster (randomly sampled) and four clusters per plant were used for each sample. Gene expression in cv. 'Cabernet Sauvignon' berries is shown relative to UBIQUITIN1 expression. Values and standard deviations derived from one qPCR run with duplicate PCR reactions on each of the four biological replicates. Further details on experimental designs and sampling methods are described in Loyola et al. (2016); Czemmel et al., (2017) and Matus et al., (2019). Asterisks indicate significant differences ($p<0.05$) between treatments based on two-way ANOVA followed by Tukey's post hoc test.

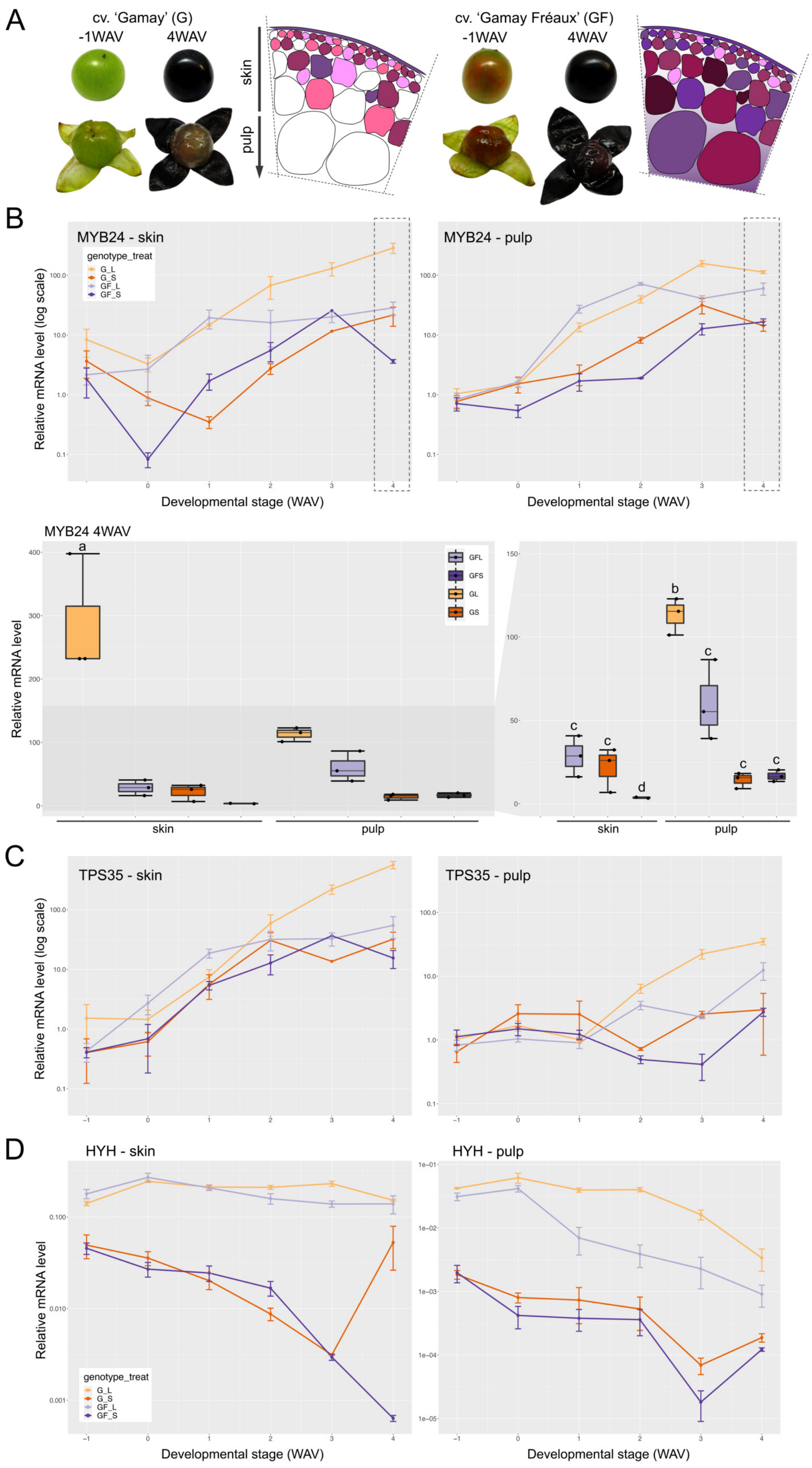


Figure S16: legend (next page)

Figure S16 (previous page): Light responsiveness of the MYB24, TPS35 and HYH in grape berry tissues. Skin and mesocarp gene expression responses to sunlight exclusion in field trials of cv. 'Gamay' and its teinturier (red-flesh) somatic variant cv. 'Gamay Fréaux'. A fruit sunlight exclusion treatment (0%) was imposed by covering grape clusters with opaque boxes (from two weeks before veraison to maturity) and compared to grape clusters exposed to natural light conditions as control (100% light incidence). The experimental design consisted in three blocks with three vines each (biological replicates n = 3). Three berries per cluster (randomly sampled) and three clusters per block were used for each sample. Gene expression values are the mean of three biological replicates with bars representing standard errors. Further details on experimental designs and sampling methods are described in Guan et al. (2016). Significant differences between treatments were calculated for 4WAV based on one- and two-way ANOVA followed by Tukey's post hoc test (also see Supplementary Figure S14).

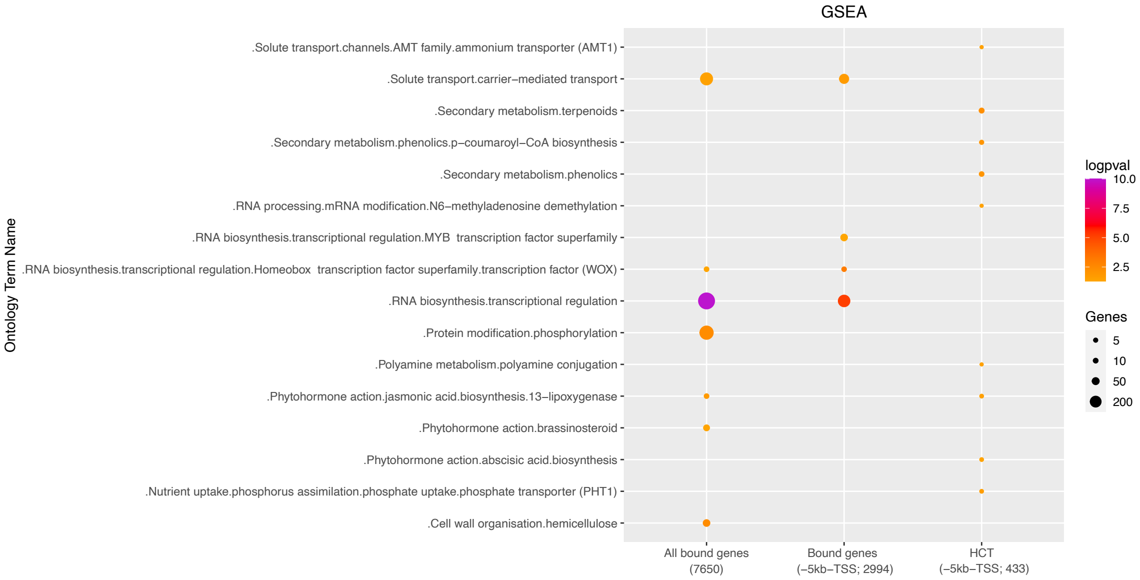


Figure S17: Figure S17: A selection of enriched terms from the MapMan gene set enrichment analysis done for i) MYB24-bound genes (DAP-seq), ii) bound genes (-5kb-TSS), and iii) High confidence target genes (-5k-TSS). A -logpval scale is provided where a higher value represents a greater statistical significance on a continuous colour scale from orange to purple. The number of genes intersecting with each term is represented by point size. Complete gene set enriched terms can be found in Supplementary Table S3.

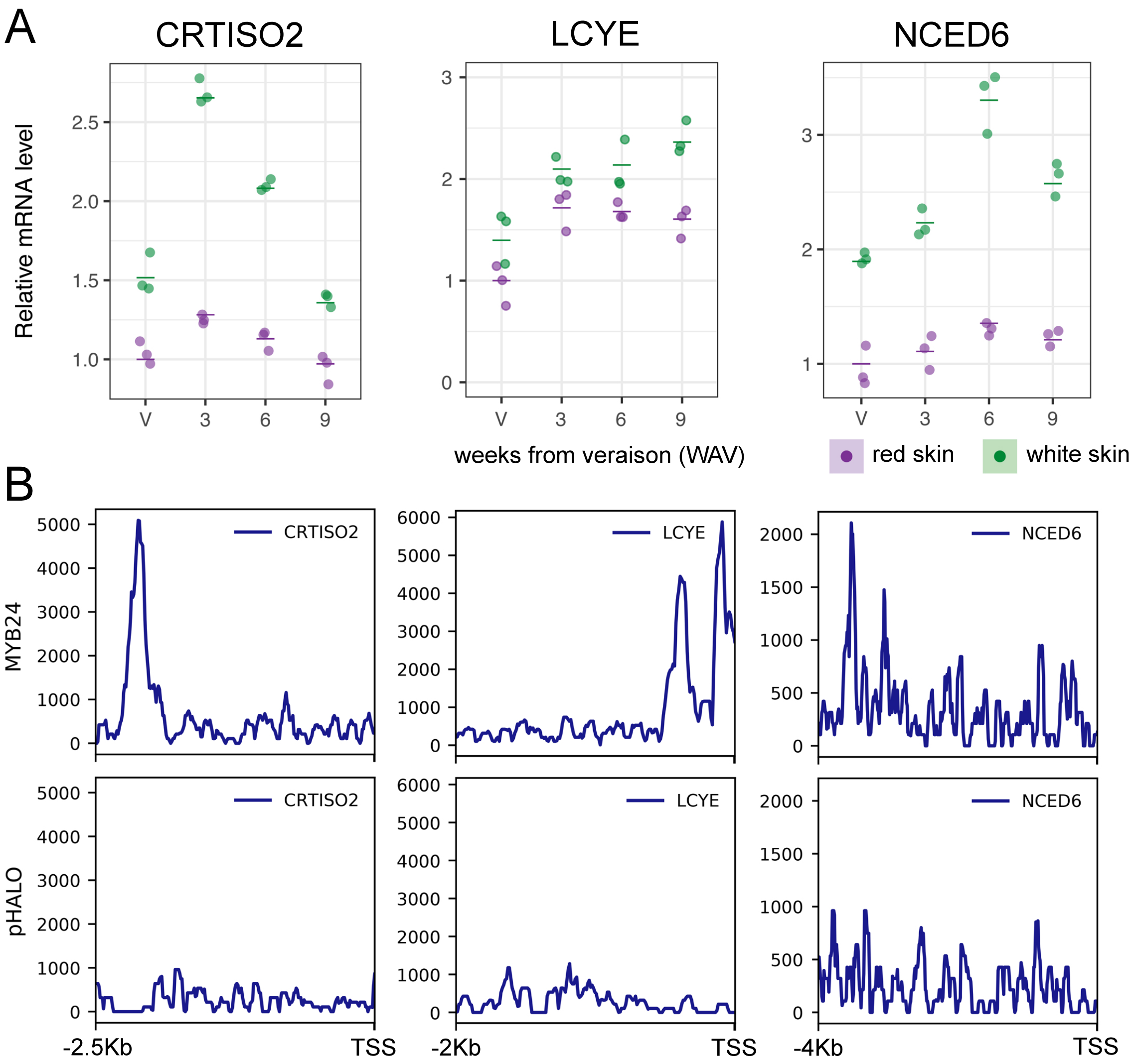


Figure S18: (A) Expression patterns of carotenoid-related genes CRTISO2 (carotenoid isomerase 2), LCYE (lycopene epsilon cyclase), and NCED6 (9-cis-epoxycarotenoid dioxygenase 6) at different ripening stages in red and white skin sections of variegated berries. About 16 berries from 8 clusters belonging to 5 plants were used per sample. Data from three biological replicates were shown (averages as horizontal lines). (B) MYB24 binds to the promoter of carotenoid-related genes (CRTISO2, LCYE, and NCED6). DAP-seq binding signal at -2.069kb (CRTISO2), -0.134kb and -0.406kb (LCYE), and -3.703kb (NCED6) from the TSS (x axis), compared to empty vector (pIX-HALO) as negative control.
